# Asymmetric activation of microglia in the hippocampus drives anxiodepressive consequences of trigeminal neuralgia

**DOI:** 10.1101/2022.04.16.488241

**Authors:** Li-Qiang Chen, Xue-Jing Lv, Qing-Huan Guo, Su-Su Lv, Ning Lv, Jin Yu, Wen-Dong Xu, Yu-Qiu Zhang

**Affiliations:** State Key Laboratory of Medical Neurobiology and MOE Frontiers Center for Brain Science, Department of Translational Neuroscience, Jing’an District Centre Hospital of Shanghai, Institutes of Brain Science, Fudan University, Shanghai 200032, China; Department of Integrative Medicine and Neurobiology, State Key Laboratory of Medical Neurobiology, School of Basic Medical Sciences, Shanghai Medical College, Fudan University, Shanghai, 200032, China; Department of Hand Surgery, Huashan Hospital, Fudan University, Shanghai 200040, China

**Author notes:** Correspondence: Yu-Qiu Zhang Ph.D. Institutes of Brain Science, Fudan University #B5-005 Scientific Research Building B, 131 Dong An Road, Shanghai 200032, China, (Y.-Q. Z.). All of the authors have no competing financial interest in this study.

**Keywords:** Trigeminal neuralgia, Microglial activation, Anxiodepression, Hippocampus, long-term potentiation (LTP)

## Abstract

Patients suffering from trigeminal neuralgia (TN) are often accompanied by anxiety and depression. Whether and how microglia are involved in TN-induced anxiodepressive remains unclear. Here, we unconventionally report that TN activates ipsilateral but not contralateral hippocampal microglia, upregulates ipsilateral hippocampal ATP and interleukin1β (IL-1β) levels, impairs ipsilateral hippocampal long-term potentiation (LTP), and induces anxiodepressive-like behaviors in a time-dependent manner in rodents. Specifically, activation of ipsilateral hippocampal microglia is necessary for TN-induced anxiodepressive-like behaviors; and unilateral activating hippocampal microglia is sufficient to elicit an anxiodepressive state and impair LTP. Knockdown of ipsilateral hippocampal P2X7 receptor prevented TN-induced microglial activation and anxiodepressive-like behaviors. Furthermore, we demonstrate that microglia-derived IL-1β mediates microglial activation-induced anxiodepressive-like behaviors and LTP impairment. Together, these findings suggest that priming of microglia with ATP/P2X7R in the ipsilateral hippocampus drives pain-related anxiodepressive-like behaviors via IL-1β. Our results also reveal an asymmetric role of the bilateral hippocampus in TN-induced anxiety and depression.

## Introduction

Chronic neuropathic pain has become a leading cause of disability, and as a persistent stressor results in multiple neuropsychiatric disorders, such as anxiety and depression (1, 2). Clinically, depressions are frequently observed in patients suffering from chronic pain, with a prevalence rate of 30% in neuropathic pain patients (3), and this consequence of pain can be preclinically modeled (4–7). Patients with chronic pain and depression are poorly responsive to current antidepressant and analgesics treatment, which adds dramatically to the patients’ burden on health care services (8). Although the co-existence of chronic pain and depression has long been recognized clinically, mechanism-based preclinical studies are still rare.

In past decade, accumulating evidence has shown that microglia-mediated neuroinflammation is involved in the development of affective disorders including pathogenesis of depression (2, 9, 10). Microglia are the primary immune cells in the central nervous system and function as active surveyors of the extracellular environment in both healthy and disordered brain (11, 12). They can be activated by various extracellular stimuli and promote inflammatory responses by expressing and releasing proinflammatory cytokines, which as powerful neuromodulators regulate synaptic transmission and plasticity (13, 14). Depressed patients exhibited increased inflammatory cytokines, such as interleukin-1β (IL-1 β) and interferon-γ (IFN-γ) in several brain areas, including the hippocampus (15, 16). Imaging and postmortem analysis of depressive patients, especially suicide patients with depression, showed a significant activation of microglia in several emotion-related brain areas, such as prefrontal cortex, anterior cingulate cortex, and hippocampus (17, 18). Similar increases in microglial activation in chronic pain patients (19) and animal models (20, 21) were observed in emotion-related brain regions including hippocampus, a brain region that is strongly linked with depression and chronic pain (22, 23). The anti-depression effects of microglia activity inhibition were demonstrated in both human and in animals (24, 25), suggesting an important role of microglial activation in the pathogenesis of depression. However, there are few reports regarding the mechanism of microglia in anxiodepressive consequences of chronic pain.

Here we will reveal new insights into the role of microglia-to-neurons communication in neuropathic pain-induced depression. We used optogenetic approach in CX3CR1::ChR2 and CX3CR1::Arch mice to investigate the role of microglial activity for anxiodepressive state and long-term potentiation (LTP) of neuronal activity in the hippocampal CA1 area. We found that hippocampal microglial activation is necessary for the anxiety and depression induced by neuropathic pain, and activating microglial is sufficient to induce an anxiodepressive state. Moreover, we also show that hippocampal microglia P2X7 receptor triggers upregulation of the proinflammatory cytokine IL-1β to impair hippocampal LTP and drive depressive-like behaviors. In addition, a laterality effect has been reported in association of hippocampal volume and function with depression (26) and pain (27), indicating a specialized role for hippocampal subfields. In the present study, we provide sound evidence that ipsilateral but not contralateral hippocampal microglia are involved in depressive consequences of neuropathic pain.

## Results

### Activation of microglia in the ipsilateral hippocampal CA1 area contributes to trigeminal neuralgia-induced anxiodepressive-like behaviors in Wistar rats

Trigeminal neuralgia rat model was established by constriction of the infraorbital nerve (CION, Figure 1A) to mimic clinical trigeminal neuropathic pain. Following CION, mechanical allodynia developed with 4 days and persisted for at least 20 days in the ipsilateral, but not contralateral, vibrissa pad (2-way repeated measured ANOVA, treatment: F_2,25_=29.16, p<0.0001; treatment×time: F_6,75_=2.92, p=0.016; Figure 1B and C; Supplemental Figure 1A). We also observed significant cold allodynia at days 10 and 20 in the ipsilateral vibrissa pad of CION rats (Supplemental Figure 1B and C).

**Figure 1.**
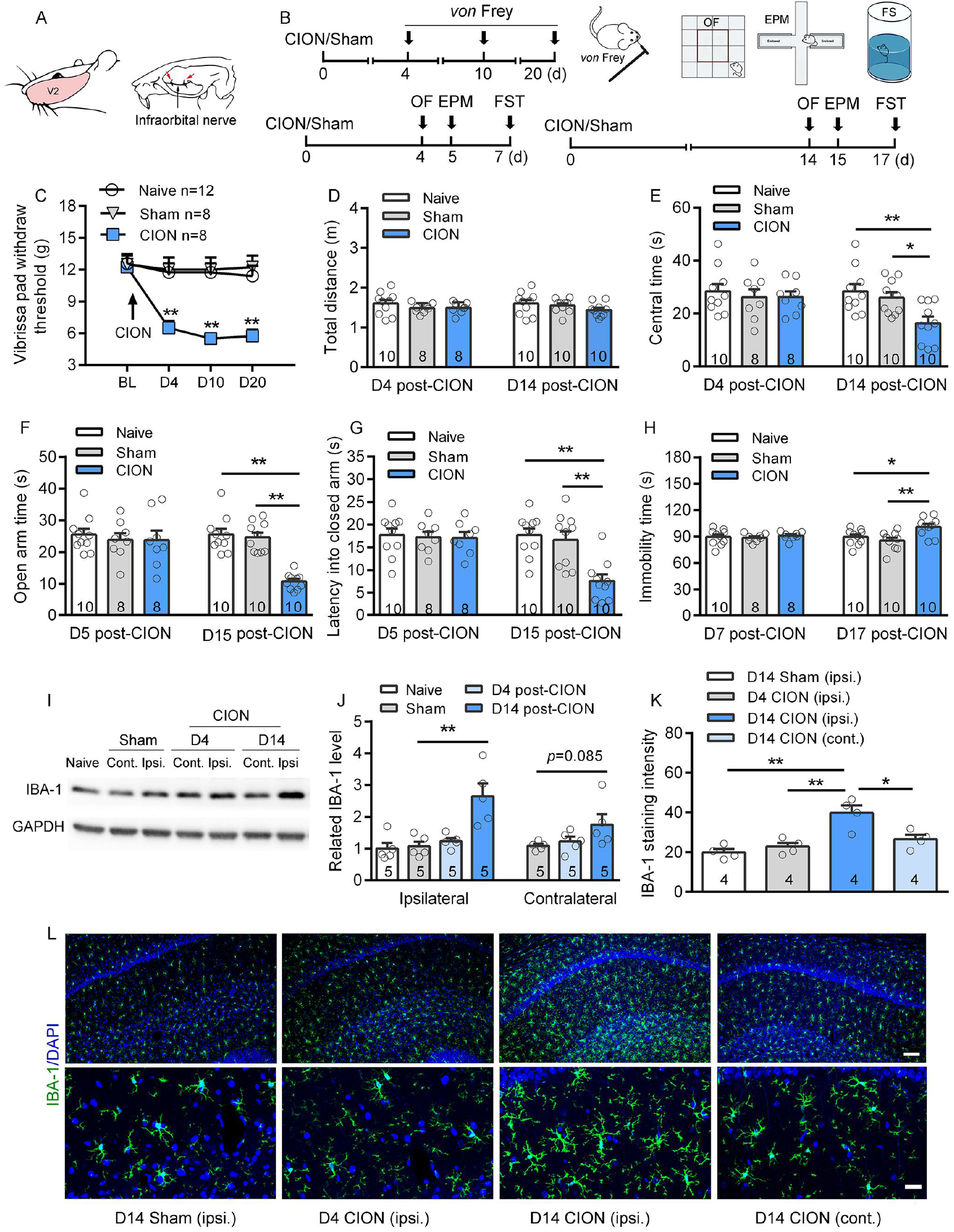
Trigeminal neuralgia time-dependently induces anxiodepressive-like behaviors, mechanical allodynia of the ipsilateral vibrissa pad and microglial activation in ipsilateral hippocampal CA1. (A) Schematic showing chronic constriction injury of infraorbital nerve (CION). Arrows point to the ligation sites. (B) Schematic of the protocol for the experiments C-H. (C) Following the trigeminal neuralgia (TN), mechanical response threshold of the ipsilateral vibrissa pad decreases markedly on day 4 and lasted for more than 20 days. \*\**P*<0.01 versus sham and naïve controls, 2-way RM ANOVA followed by post hoc Student-Newman-Keuls test (n=12/8/8, naïve/ sham/ CION, rats/group). (D-H) The TN rats exhibit anxiety-like behaviors in open field test (OF, D and E) and elevated plus maze (EPM) test (F and G), and depressive-like behavior in forced swimming test (FS, H) on days 14-17 but not days 4-7 after CION. \**P*<0.05, \*\**P*<0.01, 1-way ANOVA followed by post hoc Student-Newman-Keuls test (n=10/8/8, naive/sham-/CION-4/5/7d; n=10/10/10, naive/sham-/CION-14/15/17d, rats/group). (I and J) Western blot analysis reveals significant upregulation of IBA-1 (microglial marker) level on day 14 after CION in the ipsilateral hippocampal CA1 area. \*\**P*<0.01, 1-way ANOVA followed by post hoc Student-Newman-Keuls test (n=5 rats for all the groups). (K and L) Immunohistochemistry data shows microglial activation, as indicated by intense IBA-1 immunofluorescence and large cell bodies of microglia in the ipsilateral hippocampal CA1 area. Microglial activation occurs on day 14 but not day 4 after CION. \**P*<0.05, \*\**P*<0.01, 1-way ANOVA followed by post hoc Student-Newman-Keuls test (n=4 rats for all the groups). Scale bars indicate 100 μm and 25 μm for upper and bottom rows of l, respectively.

The anxiodepressive-like behavioral tests were performed in different groups from day 4 to day 7 and day 14 to day 17, respectively (Figure 1B). In the anxiety-like behavioral tests, the rats with CION for more than 14 days but not for 4/5 days spent less time in the center arena of open field (OF, 1-way ANOVA, F_2,27_=6.87, p=0.0039, Figure 1D and E; Supplemental Figure 1D) and in the open arm of elevated plus maze (EPM, 1-way ANOVA, F_2,27_=33.44, p<0.0001), as well as shorter latency into the closed arm in EPM (1-way ANOVA, F_2,27_=12.2, p=0.0002, Figure 1F and G; Supplemental Figure 1D). Similarly, CION rats also exhibited depressive-like behavior at day 17 but not day 7 after surgery in the forced swimming (FS, Figure 1H) test. The rats with CION for more than 2 weeks exhibited more immobile duration in FS test relative to the naive, sham and CION rats for 1 week (1-way ANOVA, F_2,27_=6.64, p=0.0045; Figure 1H). These results indicated that anxiodepressive-like behaviors caused by neuropathic pain were time-dependent. There was no difference among the groups for total travel distance in OF (Figure 1D) and latency to fall in rotarod test (Supplemental Figure 1E), indicating that the above behavioral phenotypic differences were not due to motor impairment.

It has been reported that microglia and neuroinflammtion in the brain, such as anterior cingulate cortex (28) and dorsal striatum (10), contribute to the pathogenesis of anxiety and depression. In the present study, we further examined whether hippocampal microglia were involved in CION-induced anxiety and depression. Western blot analysis showed a robust elevation of IBA-1 (a microglial marker) in the ipsilateral hippocampal CA1 area on day 14 but not day 4 after CION, which was associated with the development of anxiodepressive-like behaviors (1-way ANOVA, F_3,16_=10.89, p=0.0004; Figure 1I and J). CION-induced activation of microglia in the ipsilateral hippocampus after 14 days was further confirmed by immunohistochemistry (Figure 1K and L). Activated microglia exhibited large cell bodies and short or thick processes in the rat hippocampal CA1 area (Figure 2A-D). Consistently, CD68 (a microglial activation marker) expression in the hippocampal CA1 area were also significantly increased at day 14 after CION (Figure 2E and F).

**Figure 2.**
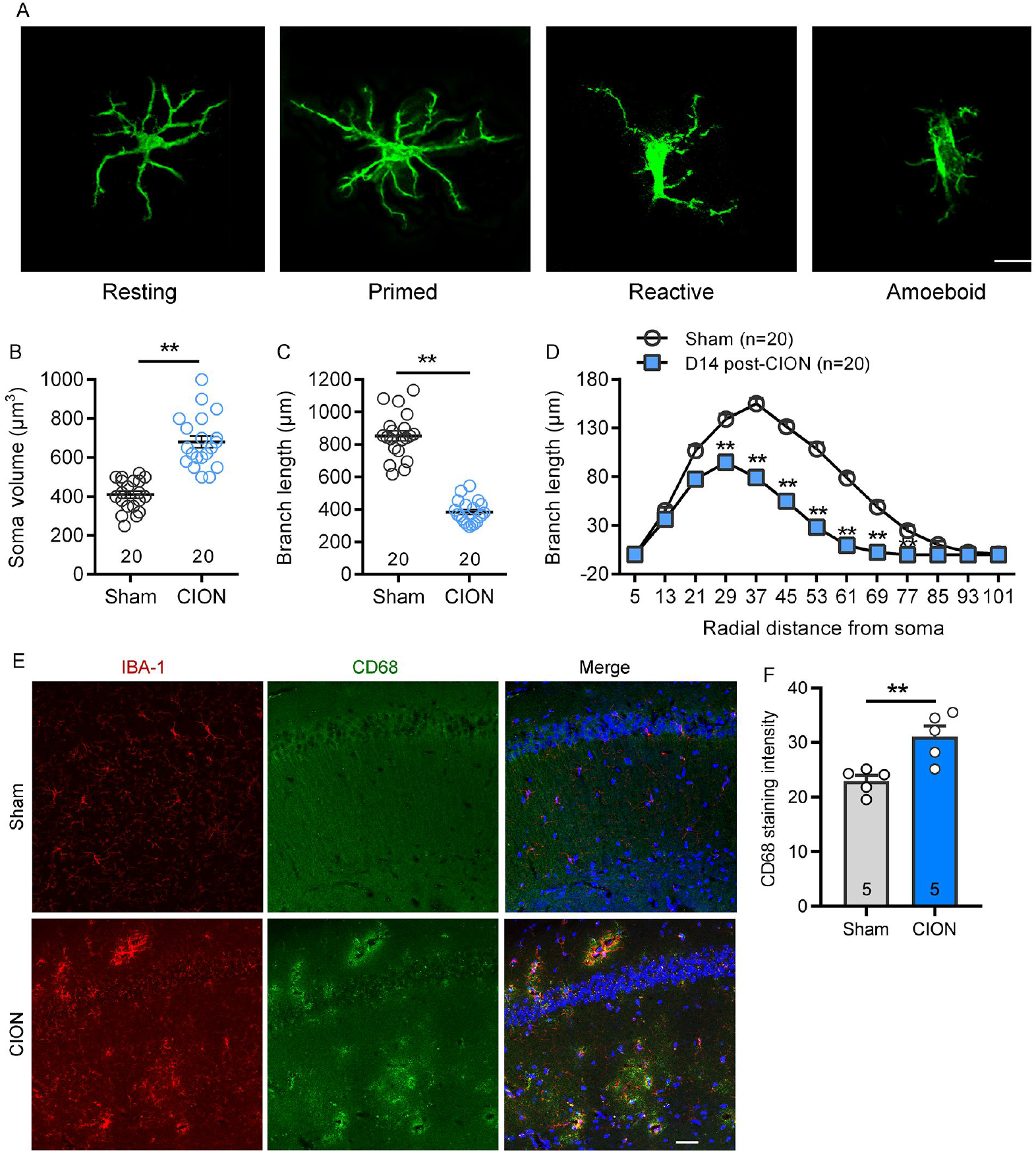
Morphological changes and activation of hippocampal microglia at 14 day after CION. (A) Representative immunofluorescence images showing the morphology of microglia in resting (radial branching), primed (thickened processes, increased polarity), reactive (thickened stout processes with highly reduced branching), and amoeboid (few-to-no processes, enlarged cell body). Scale bar indicates 20 μm. (B-D) Sholl analysis showing enlarged soma size (B) and shortened branches (C and D) of hippocampal microglia on day 14 after CION. \*\**P* < 0.01 versus sham, two-tailed Student’s *t* test (B and C) or 2-way RM ANOVA followed by post hoc Student-Newman-Keuls test (D, n=20 cells for sham and CION-14 day). (E and F) Double immunofluorescence staining of CD68 and IBA-1 showing CD68-IR coexpressed with IBA-1, and increased CD68 positive signal in the hippocampal CA1 area on day 14 after CION in mice. Scale bars indicate 30 μm. \*\**P* < 0.01, two-tailed Student’s *t* test (n=5/5, sham/CION, mice/group).

Next, we examined the effects of temporary partial deletion hippocampal microglia on development of CION-induced anxiodepressive-like behaviors in rats. We microinjected Mac-1-SAP (2.5 μg/1 μl, per side), a specific microglia cytotoxin targeting CD11b expressing cells (29, 30), into the hippocampal CA1 area on day 0 and day 7 after CION (Figure 3A and B), a robust decrease in the IBA-1-immunoreactivity (IBA-1-IR) was observed at least for 2 weeks. Following Mac-1-SAP into the bilateral hippocampal CA1 area, CION-induced IBA-1 upregulation was significantly suppressed (Figure 3C and D), and the CION rats failed to develop anxiodepressive-like behaviors in OF test (1-way ANOVA, central time: F_5, 30_=7.36, p=0.0001; central distance percentage: F_5, 30_=9.46, p<0.0001; Figure 3E and F), EPM test (1-way ANOVA, open arm time: F_5, 30_=6.72, p=0.0003; latency into closed arm: F_5, 30_=8.29, p<0.0001; Figure 3G and H) and FS test (1-way ANOVA, F_5, 30_=11.89, p<0.0001; Figure 3I), suggesting hippocampal microglia play an important role in chronic pain-induced anxiety and depression. The anti-anxiodepressive effect of inhibiting microglia was further confirmed by systematic administration of minocycline (20 mg/kg, i.p.), a semisynthetic second-generation tetracycline that has emerged as a potent inhibitor of microglial activation (Supplemental Figure 2).

**Figure 3.**
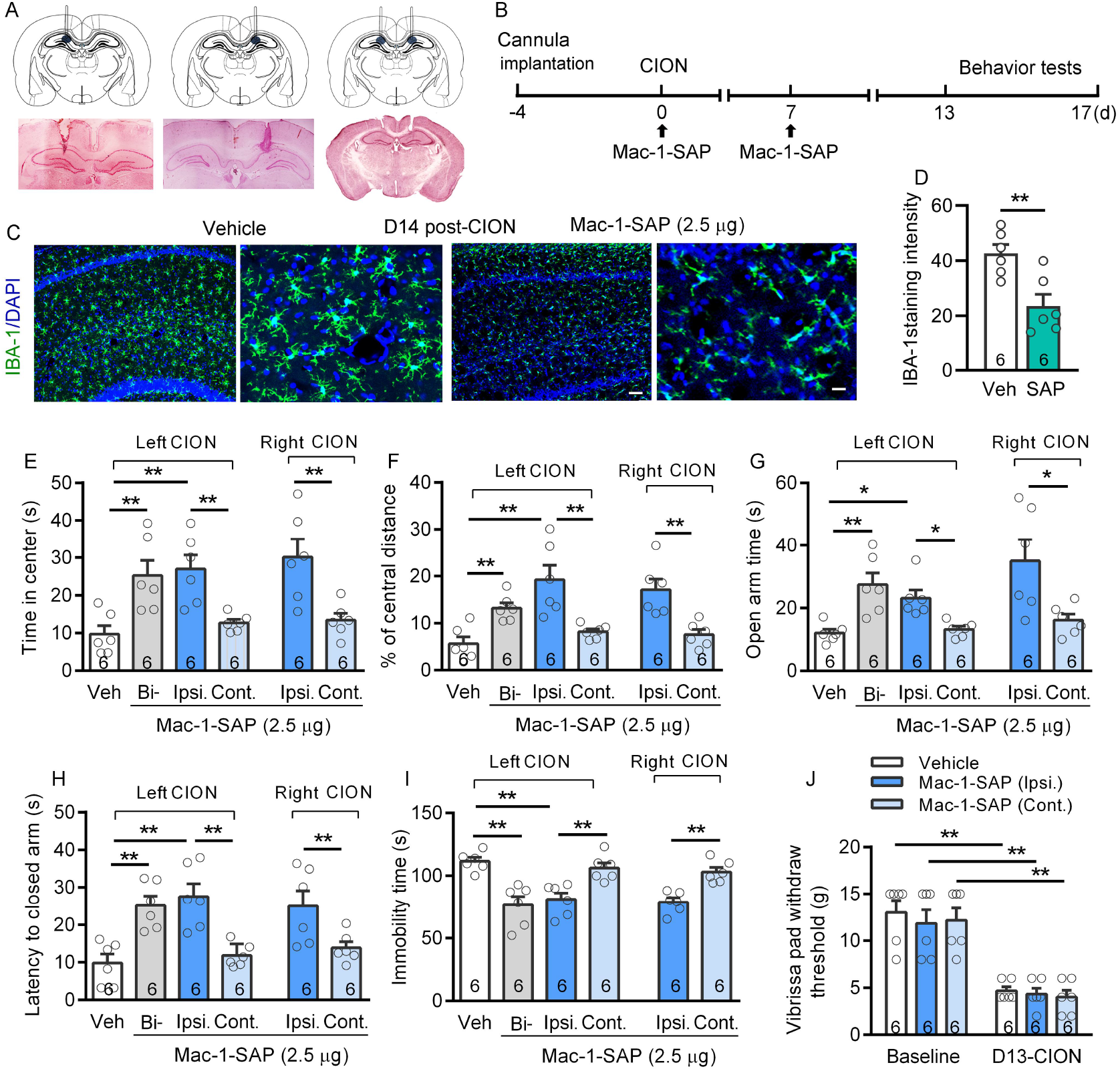
Temporary depletion of hippocampal microglia blocks the CION-induced anxiodepressive-like behaviors. (A) Schematic and photomicrograph of coronal section showing cannula placement in the unilateral and bilateral hippocampus. (B) Schematic of the protocol for the experiments E-J. (C and D) Selective targeting of hippocampal microglia with Mac-1-SAP (2.5 μ g, once a week for twice) results in a robust decrease in the immunofluorescence staining signal of IBA-1 on day 14. Scale bars indicate 100 μm (low magnification) and 25 μm (high magnification). \*\**P*<0.01, two-tailed Student’s t test (n=6 rats or all the groups). (E-I) Ipsilateral or bilateral injection of Mac-1-SAP (2.5 μg, twice) into the CA1 area of hippocampus leads to a significant anti-anxiodepressive effect in OF (E and F), EPM (G and H) and FS (I) tests in CION rats. Administration of Mac-1-SAP into the contralateral hippocampal CA1 fails to block the development of anxiodepressive-like behaviors, no matter the CION on the left or right. \**P*<0.05, \*\**P*<0.01, 1-way ANOVA followed by post hoc Student-Newman-Keuls test (n=6 rats for all the groups). (J) Neither ipsilateral nor contralateral intra-CA1 of Mac-1-SAP affects CION-induced mechanical allodynia. \*\**P*<0.01, 1-way ANOVA followed by post hoc Student-Newman-Keuls test (n=6 rats for all the groups).

Intriguingly, administration of Mac-1-SAP to the left (ipsilateral to CION) but not right (contralateral to CION) hippocampus can mimic the anxiolytic and antidepressant effect of bilateral Mac-1-SAP. To further confirm that ipsilateral rather than left hippocampal microglia are involved in the development of CION-induced anxiodepressive-like behaviors, we performed right CION surgery and compared the effects of Mac-1-SAP given to the ipsilateral (right) and contralateral (left) hippocampal CA1 area. As shown in Figure 3E-I, partial deletion of ipsilateral hippocampal microglia prevented the development of anxiodepressive-like behaviors by neuropathic pain, regardless of left or right CION. Differently, neither ipsilateral nor contralateral administration of Mac-1-SAP alleviated CION-induced mechanical allodynia (Figure 3J). These results indicate that ipsilateral hippocampal microglia are required for the development of neuropathic pain-induced anxiodepressive-like behaviors in trigeminal injury rats.

### Optogenetic manipulation of ipsilateral hippocampal microglia in the CA1 area alters the anxiodepressive state in mice

Just like Wistar rats, C57/BL6 mice with CION exhibited similar anxiodepressive-like behaviors after 2 weeks but not within one week (Supplemental Figure 3). To obtain selective control of microglial activity, we crossed CX3CR1-CreER mice, harboring a tamoxifen-inducible Cre recombinase, with Ai35 mice carrying the floxed stop-Arch-EGFP gene in the ROSA26 locus, to generate the CX3CR1::Arch mice. The optic fiber was implanted in the CA1 area of hippocampus ipsilateral to CION and yellow light (580 nm) illumination was applied from days 7 to 15 after CION (Figure 4A and B). As shown in Figure 4C, Arch-EGFP signals were colocalized with IBA-1-IR after tamoxifen injection. Following yellow light-treatment, CION-induced microglial activation of the hippocampal CA1 area was suppressed in CX3CR1::Arch mice (Figure 4D and E). Consistent with the inhibition of microglia by Mac-1-SAP into ipsilateral CA1, optogenetic inhibition of ipsilateral hippocampal microglia significantly blocked CION-induced anxiodepressive-like behaviors in OF, EPM and FS tests (Figure 4F-J).

**Figure 4.**
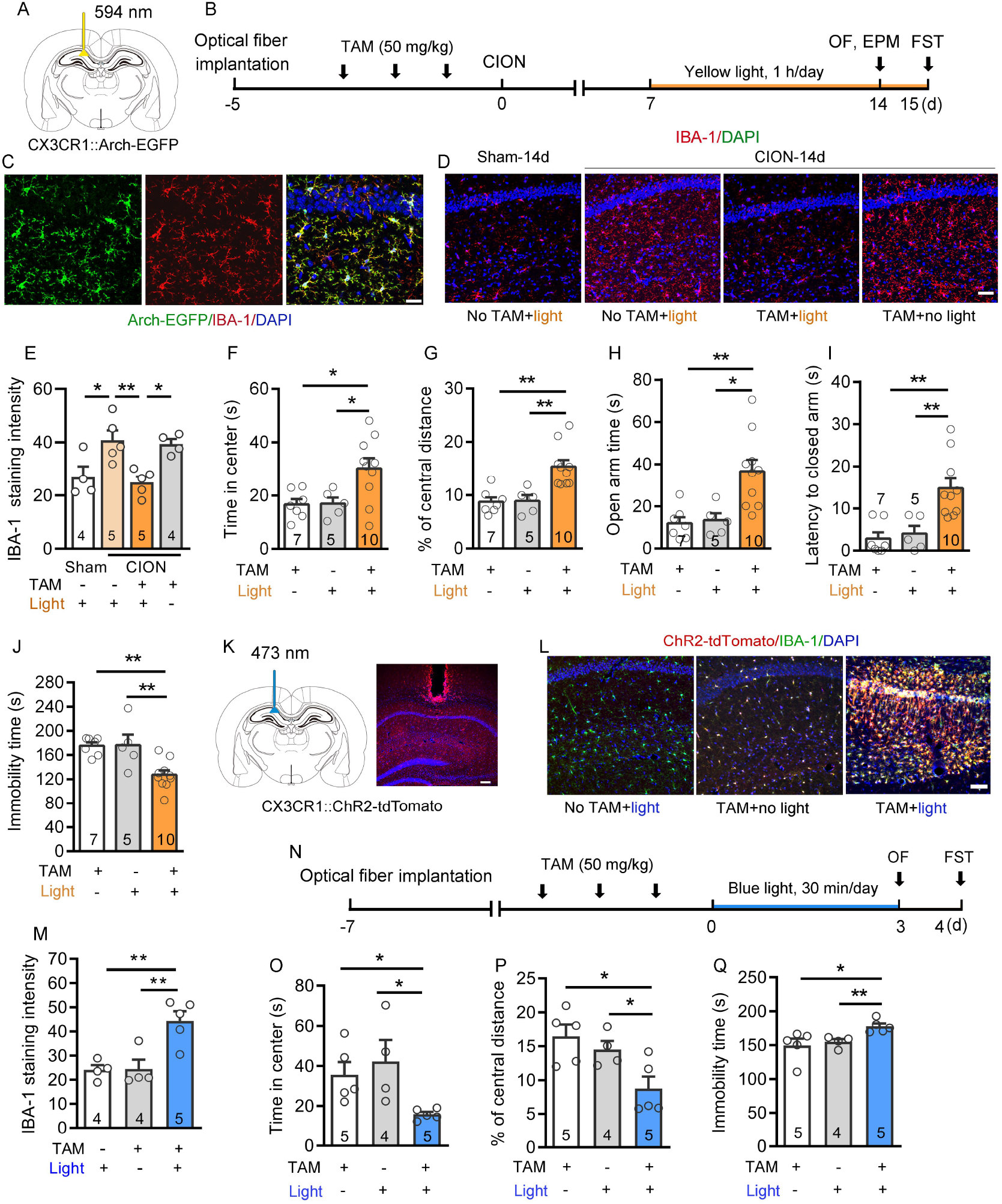
Optogenetic manipulation of unilateral hippocampal microglia in the CA1 area alters the anxiodepressive-like behaviors in mice. (A) Schematic showing the site of optical fiber implantation in the unilateral hippocampal CA1 area of CXCR1::Arch mouse. (B) Schematic of the protocol for experiments F-J. (C) Double immunofluorescence staining reveals that Arch-EGFP coexpressed with IBA1. Scale bar: 50 μm. (D and E) IBA-1 immunoreactivity in the hippocampal CA1 area is reduced by yellow light (580 nm, 50 ms on, 10 ms off) illumination on day 14 after CION in CXCR1::Arch mice. Scale bar: 50 μm. \**P*<0.05, \*\**P*<0.01, 1-way ANOVA followed by post hoc Student-Newman-Keuls test (n=4/5/5/4, sham+light/CION+light without TAM/CION+TAM+light/CION+TAM without light, mice/group). (F-J) Optogenetic inhibition of ipsilateral hippocampal microglia prevents the anxiodepressive-like behaviors in OF (F and G), EPM (H and I) and FS (J) tests on day 14/15 after CION in CXCR1::Arch mice. \**P*<0.05, \*\**P*<0.01, 1-way ANOVA followed by post hoc Student-Newman-Keuls test (n=7/5/10, no light/no TAM/TAM+light, mice/group)). (K) Schematic and photomicrograph of coronal section showing the site of optical fiber implantation in the unilateral hippocampal CA1 area of CXCR1::ChR2 mouse. Scale bar: 100 μ m. (L and M) IBA-1 immunoreactivityin the hippocampal CA1 area is increased by blue light (473 nm, 20 HZ, 25 ms) illumination on day 14 after CION in CXCR1::ChR2 mice. Scale bar: 50 μ m. \*\**P*<0.01, 1-way ANOVA followed by post hoc Student-Newman-Keuls test (n=4/4/5, no TAM/no light/TAM+light, mice/group). (N) Schematic of the protocol for experiments O-Q. (O-Q) Optogenetic activation of unilateral hippocampal microglia induces anxiodepressive-like behaviors in OF (O and P) and FS (Q) tests in naive CX3CR1::ChR2 mice. \**P*<0.05, \*\**P*<0.01, 1-way ANOVA followed by post hoc Student-Newman-Keuls test. (n=5/4/5, no light/no TAM/TAM+light, mice/group).

We also examine the effects of activating hippocampal microglia on anxiety and depression levels in normal mice. We crossed CX3CR1-CreER line with floxed ChR2-tdTomato mice to generate the CX3CR1::ChR2 mice. Implantation of optic fiber per se did not cause obviously microglial activation in the CA1 area of hippocampus (Figure 4K). ChR2-tdTomato signals were colocalized with IBA-1-IR after tamoxifen injection, and blue light (473 nm) illumination evoked a robust activation of microglia in the hippocampal CA1 area of CX3CR1::ChR2 mice (Figure 4L and M). Optogenetic activation of unilateral hippocampal microglia elicited an obvious anxiodepressive-like behavioral phenotype, marked by less center time (1-way ANOVA, F_2, 11_=4.31, p=0.042) and distance (1-way ANOVA, F_2, 11_=6.15, p=0.016) in OF test (Figure 4O and P), and higher immobile duration in FS test (1-way ANOVA, F_2, 11_=4.66, p=0.034; Figure 4Q) than that of controls. Taken together, the above results suggest that unilateral hippocampal CA1 microglial activation is sufficient to drives mouse anxiodepressive-liker behaviors.

### Trigeminal neuralgia impairs ipsilateral hippocampal LTP by activated microglia in mice

Impaired hippocampal long term potentiation (LTP) has been found in a variety of animal models of depression (31–34). To investigate whether hippocampal LTP is affected by trigeminal neuralgia and related anxiodepression, we recorded hippocampal CA1 area LTP in naive, sham and post-CION 4 day and 14 day mice. Stimulation of Schaffer collaterals evoked a basal field excitatory postsynaptic potential (fEPSP) in the CA1 area and 4 trains of theta burst stimulation (TBS) induced LTP, which was maintained for at least 60 min. LTP was successfully induced in naive, sham and post-CION 4 day mice (Figure 5A and 4B). However, by 14 days after CION, ipsilateral hippocampal LTP was impaired, with only short-term potentiation, regardless of the left or right CION (Figure 5B and C). The functional output of microglia has been proposed to produce via neuronal activity (35). Most evidence showed that microglia regulate synaptic plasticity and respond to multiple behavioral consequences, such as depression and chronic pain (9, 12). To investigate whether the suppression of hippocampal LTP in ipsilateral to CION can be rescued by inhibition of microglia, we recorded LTP in hippocampal slices from the ipsilateral side of CION in the presence of minocycline (5 μmol/L) or vehicle (Figure 5D). The suppression of hippocampal LTP was completely rescued by minocycline treatment (Figure 5E). We also recorded hippocampal LTP in CX3CR1::Arch mice. Yellow light (580 nm) illumination was applied from days 7 to 14 after CION with tamoxifen or without tamoxifen (control) treatment mice (Figure 5F). Optogenetic inhibition of microglia in the ipsilateral hippocampal CA1 area significantly blocked CION-induced LTP suppression (Figure 5G), Furthermore, we examine the effects of directly activating microglia on hippocampal LTP by optogenetically stimulating the hippocampal slices from CX3CR1::ChR2 mice (Figure 5H). Blue light (473 nm, 20 HZ, 25 ms) illumination suppressed hippocampal LTP, with a decreased fEPSP amplitude as compared to no light control and no tamoxifen control groups (Figure 5I). These results suggest that microglial activation in the hippocampal CA1 area may be involved in mediating the impairment of LTP.

**Figure 5.**
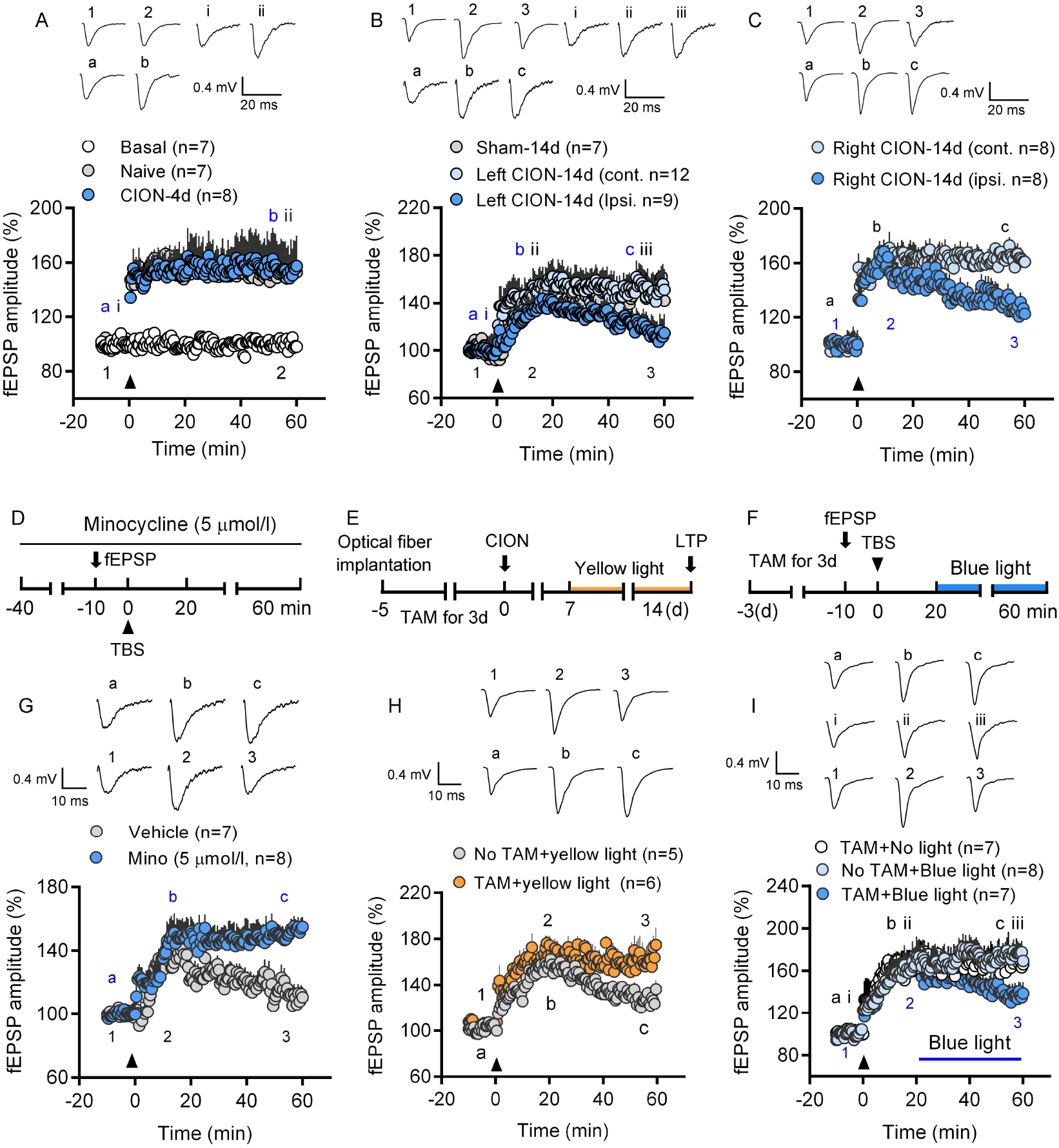
Trigeminal neuralgia time-dependently impairs ipsilateral hippocampal long-term potentiation by activated microglia. (A) Long-term potentiation (LTP) of fEPSP in area CA1 of hippocampus is successfully induced by the TBS protocol (indicated by an arrow head) in hippocampal slice from both naive and CION-4 day mice (n=7/7/8, basal/naive/CION-4 day, slices/group). Stable basal recording (without TBS) suggested a stable electric stimulation. (B and C) The LTP in the ipsilateral hippocampal CA1 area is impaired on day 14, regardless of the left (B) or right (C) CION (n=7/12/9, sham/contralateral/ipsilateral hippocampal in left CION on day 14; n=8/8, contralateral/ipsilateral hippocampal in right CION on day 14, slices/group). (D and E) Pre-application of minocycline (5 μmol/L) reverses CION-induced LTP impairment (n=7/8, vehicle/minocycline, slices/group). (F and G) Optogenetic inhibition of hippocampal microglia partially blocks CION-induced ipsilateral hippocampal LTP impairment in CX3CR1::Arch mice (n=5/6, no TAM+yellow light/TAM+yellow light, slices/group). (H and I) Optogenetic activation of hippocampal microglia suppressed LTP amplitude in CX3CR1::ChR2 mice (n=7/8/7, TAM+no blue light/no TAM+blue light/TAM+blue light, slices/group). The insets showing the traces of fEPSPs before and after TBS stimulation in different treatment groups (point 1 and 2, point i and ii, or point a, b).

### P2X7R mediates CION-induced hippocampal microglial activation and anxiodepressive-like behaviors

The previous studies from our laboratory showed that P2X7R was expressed in spinal and hippocampal microglia, and mediated bone cancer pain and chronic stress-induced depression (36, 37). In the current study, we further identified whether P2X7R mediates hippocampal microglial activation and neuropathic pain-induced anxiodepressive-like behaviors. We performed western blotting to identify the protein expression of P2X7R in the bilateral hippocampal CA1. There was a significant upregulation of P2X7R on day 14 but not day 4 after CION in the ipsilateral hippocampus (1-way ANOVA, F_6, 28_=4.18, p=0.004; Figure 6A and B), which is consistent with the temporal spatial characteristics of microglial activation. The increased P2X7R expression in the ipsilateral CA1 at 14 days after CION was further confirmed by immunohistochemistry (Figure 6C). Specificity of P2X7R antibody was verified in *p2x7* receptor gene knock-out mice by western blot and immunohistochemistry (Supplemental Figure 4A and B). Double immunostaining showed that P2X7R-IR was almost coexpressed with IBA-1 (microglial marker) but not with GFAP (astrocytic marker) and NeuN (neuronal marker) in the hippocampus (Figure 6D). To confirm whether neuropathic pain results in increased release of ATP, the ligand for P2X7R, in the hippocampus, we measured ATP concentration in extracellular dialysate of the hippocampal CA1 on day 14 after CION. Compared with the sham rats, the extracellular ATP level was significantly increased in the ipsilateral hippocampus (1-way ANOVA, F_3, 20_=8.7, p=0.0007; Figure 6E). To address whether P2X7R is involved in CION-induced hippocampal microglial activation and anxiodepressive-like behaviors, we examined the effects of P2X7R knockdown in the hippocampus. The knockdown efficiency was determined by reduced P2X7R expression after treatment with the siRNA (0.4 μg) targeted against rat P2X7R, and IBA-1 upregulation on day 14 after CION was significantly inhibited by P2X7R-siRNA (Figure 6F-I; 1-way ANOVA, P2X7R: F_2, 15_=16.77, p<0.0001; IBA-1: F_2, 15_=14.52, p=0.0003). Infusion of P2X7R-siRNA into the ipsilateral but not contralateral CA1 completely blocked CION-induced anxiodepressive-like behaviors in OF, EPM and FS tests (Figure 6K-O). Neither ipsilaterally nor contralaterally knockdown of P2X7R in the hippocampal CA1 alleviated CION-induced mechanical allodynia (Figure 6P). The behavioral effects of ipsilateral hippocampal P2X7R in CION rats were further confirmed by pharmacological approach. After intra-CA1 of 14-day A740003 (0.375 μg/h), a P2X7R specific antagonist, by osmotic pump system bilaterally or ipsilaterally but not contralaterally, CION rats failed to develop anxiodepressive-like behaviors (Supplemental Figure 5A-G). Similarly, neither ipsilaterally nor contralaterally delivering A740003 in the hippocampal CA1 alleviated CION-induced mechanical allodynia (Supplemental Figure 5H). These results suggest that P2X7R in the ipsilateral hippocampus to trigeminal neuralgia was specifically involved in trigeminal neuralgia-induced anxiety and depression.

**Figure 6.**
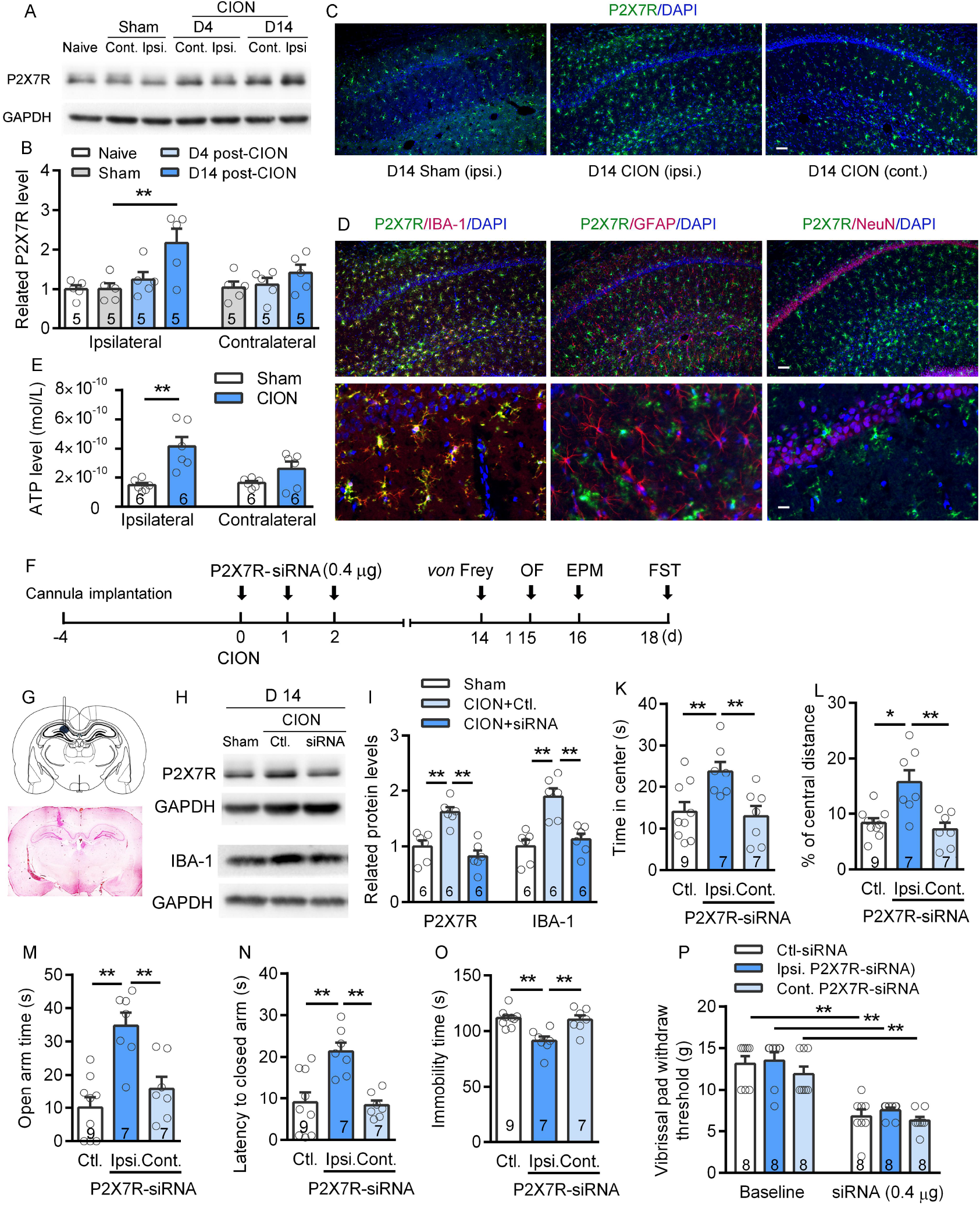
P2X7 receptor mediates CION-induced microglial activation and anxiodepressive-like behaviors. (A and B) Western blot analysis reveals significant upregulation of P2X7 receptor (P2X7R) level on day 14 after CION in the ipsilateral hippocampal CA1 area of rats. \*\**P*<0.01, 1-way ANOVA followed by post hoc Student-Newman-Keuls test (n=5 rats for all the groups). (C) Immunofluorescence staining of P2X7R showing increased PX27R positive signal in the ipsilateral hippocampal CA1 area on day 14 after CION in rats. Scale bars indicate 100 μm. (D) Double immunofluorescence staining reveals that P2X7R-IR coexpressed with IBA-1 (microglial marker) but not GFAP (astrocytic marker) and NeuN (neuronal marker) in the hippocampus. Scale bars indicate 100 μm (low magnification) and 25 μm (high magnification). (E) ATP concentration in extracellular dialysate of the ipsilateral hippocampal CA1 area is significant increased on day 14 after CION. \*\**P*<0.01, 1-way ANOVA followed by post hoc Student-Newman-Keuls test (n=6 rats for all the groups). Error bars indicate standard error of the mean. See also Figure S5. (F) Schematic of the protocol for experiments H-M. (G) Schematic and photomicrograph of coronal section showing cannula placement in the unilateral hippocampus. (H and I) Knockdown of P2X7R by siRNA (0.4 μg) against P2X7R blocks CION-induced P2X7R and IBA-1 upregulation on day 14. \*\**P*<0.01, 1-way ANOVA followed by post hoc Student-Newman-Keuls test (n=6 rats for all the groups). (K-O) Knockdown of P2X7R by siRNA in the ipsilateral but not contralateral hippocampus prevents CION-induced anxiodepressive-like behaviors in OF (K and L), EPM (M and N) and FS (O) tests (n=9/7/7, control-siRNA/ipsilateral/contralateral P2X7R-siRNA, rats/group). (P) CION-induced mechanical allodynia of vibrissa pad is not blocked by both ipsilateral and contralateral intra-CA1 of P2X7R-siRNA. \*\**P*<0.01, 1-way ANOVA followed by post hoc Student-Newman-Keuls test (n=8 rats for all the groups).

Similar to the effects of systematic administration of minocycline hydrochloride, repeated intraperitoneal injection of a selective P2X7R antagonist BBG (40 mg/kg/day) also prevented the development of anxiodepressive-like behaviors and attenuated mechanical allodynia in CION rats (Supplemental Figure 6A-I). Additionally, in *p2x7r* knockout (KO) mice, we further confirmed the role of P2X7R in pain-like hypersensitivity, hippocampal LTP and anxiodepressive-like behaviors. All the CION-induced mechanical allodynia, LTP impairment and anxiodepressive-like behaviors in wild-type mice were not developed in *p2x7r* KO mice (Supplemental Figure 7A-H).

### P2X7R contributed to CION-induced anxiodepressive-like behaviors via IL-1β

Activated microglia synthesize and release various cytokines, such as IL-1β and facilitates the development of neuropathic pain (13). Thus, we reasoned that P2X7R activation could cause IL-1β upregulation in the hippocampal CA1. Consistent with the temporal profiles of P2X7R and IBA-1 upregulation, significant elevation of IL-1β level occurred in the ipsilateral hippocampus on day 14 after CION (1-way ANOVA, F_6, 21_=4.1, p=0.007; Figure 7A and B). ELISA analysis revealed that P2X7R agonist BzATP (100 ng/ml) significantly increased LPS (100 ng/ml)-stimulated IL-1β level in hippocampal slices, which was blocked by P2X7R antagonist A740003 (1-way ANOVA, F_2, 13_=8.61, p=0.004; Figure 7C). Knockdown of P2X7R markedly suppressed the CION-induced IL-1β upregulation on day 14 (1-way ANOVA, F_2, 15_=8.82, p=0.003; Figure 7D and E). Furthermore, we tested whether the LTP impairment caused by optogenetic activation of microglia is mediated by IL-1β. The presence of IL-1Ra (IL-1β receptor antagonist, 5.7 × 10^-9^ mol/L) significantly blocked optogenetic activating microglia-induced hippocampal LTP impairment in CX3CR1::ChR2 mice (Figure 7F and G). Behaviorally, intra-CA1 of IL-1Ra (5 μg) effectively prevented the anxiodepressive-like behaviors caused by optogenetic activation of microglia (Figure 7H-K). Importantly, intra-CA1 of IL-1Ra (0.2 μg/h) by osmotic pump system for 14 days effectively prevented CION-induced anxiodepressive-like behaviors (Figure 7L-Q). CION-induced hippocampal LTP impairment was also rescued by IL-1Ra (5.7 × 10^-9^ mol/L) treatment (Figure 7R and S).

**Figure 7.**
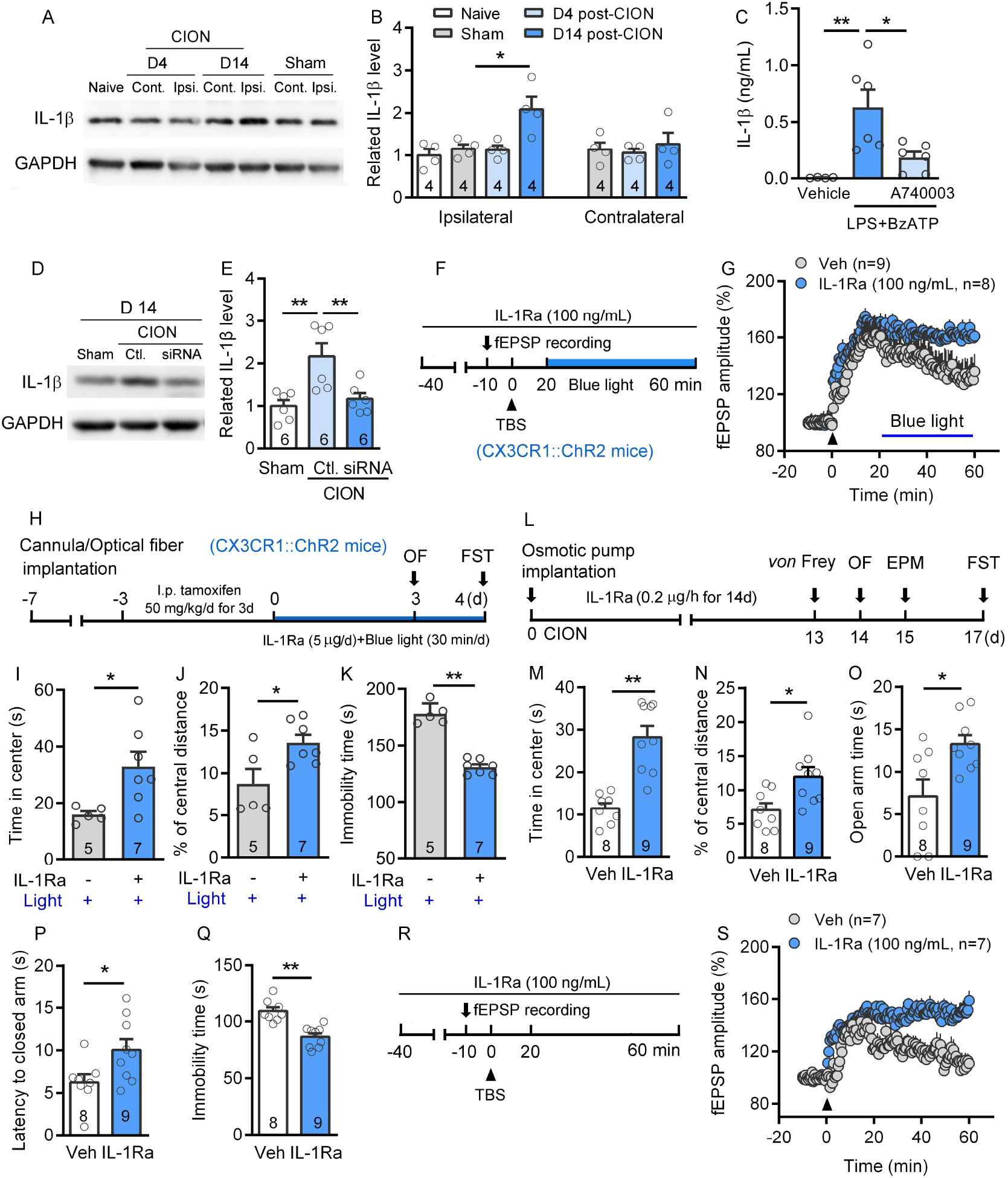
Microglial activation results in LTP impairment and anxiodepressive-like behaviors via IL-1β. (A and B) Western blot analysis reveals significant upregulation of IL-1β level on day 14 after CION in the ipsilateral hippocampal CA1 area of rats. \**P*<0.05, 1-way ANOVA followed by post hoc Student-Newman-Keuls test (n=4 rats for all the groups). (C) ELISA analysis reveals that P2X7R agonist BzATP (100 ng/ml) significantly increases LPS (100 ng/ml)-stimulated IL-1β level in hippocampal slices, and A740003 blocks the IL-1β elevation. \**P*<0.05, \*\**P*<0.01, 1-way ANOVA followed by post hoc Student-Newman-Keuls test (n=4/6/6, vehicle/A740003/no A740003, slices/group). (D and E) Western blot analysis reveals that CION-induced IL-1β upregulation is blocked by knockdown of P2X7R on day 14. \*\**P*<0.01, 1-way ANOVA followed by post hoc Student-Newman-Keuls test (n=6 rats for all the groups). (F and G) Microglial activation-induced hippocampal LTP impairment is reversed by IL-1Ra in CX3CR1::ChR2 mice (n=9/8, vehicle/IL-1Ra, slices/group). (H) Schematic of the protocol for experiments G-I. (I-M) Ipsilateral injections of IL-1Ra (5 μg/d) into the CA1 area of hippocampus blocks optogenetic activation microglia-induced anxiodepressive-like behaviors in CX3CR1::ChR2 mice. \**P*<0.05, \*\**P*<0.01, two-tailed Student’s t test (n=5/7, vehicle/IL-1Ra, mice/group). (L) Schematic of the protocol for experiments K-O. (M-Q) Intra-CA1 of IL-1Ra (an inhibitor of IL-1 receptor, 0.2 μg/h) by osmotic pump system significantly blocked CION-induced anxiodepressive-like behaviors in OF (M and N), EPM (O and P) and FS (Q) tests in CION rats. \**P*<0.05, \*\**P*<0.01, two-tailed Student’s t test (n=8/9, vehicle/IL-1Ra, rats/group). (R and S) Perfusion of IL-1Ra (5.7 × 10^-9^ mol/L) reverses CION-induced hippocampal LTP impairment in CION mice (n=7/7, vehicle/IL-1Ra, slices/group).

Next, we tested whether IL-1β per se is sufficient to impair LTP and elicit anxiodepressive-like behaviors. Acute perfusion of IL-1β (5.8 × 10^-10^ mol/L) robustly impaired the maintenance of hippocampal LTP (Figure 8A and B). Consistently, intra-CA1 of IL-1β (0.125 ng/h) by osmotic pump system directly produced anxiodepressive-like behaviors in OF, EPM and FS tests (Figure 8C-H). These results suggest that hippocampal microglial derived IL-1β contributes to CION-induced anxiety and depression. IL-1β has been reported to increase the activity of indoleamine 2,3-dioxygenase 1 (IDO1), a rate-limiting enzyme in tryptophan metabolism (38). Increased IDO resulted in the increased kynurenine (KYN)/tryptophan ratio and decreased serotonin (5-HT)/tryptophan ratio in the bilateral hippocampus (39). The resultant reduction of 5-HT may link to the monoamine hypothesis of major depression (38). Therefore, we also examined the effect of IDO1 inhibitor on CION-induced anxiodepressive-like behaviors. When 1-methyl-D-tryptophan (1-MT, 0.2 μg/h), a selective inhibitor of IDO1, was delivered into ipsilateral hippocampal CA1 area for consecutive 14 days after CION by osmotic pump system, CION rats failed to develop anxiodepressive-like behaviors in 14 days (Figure 8I-N).

**Figure 8.**
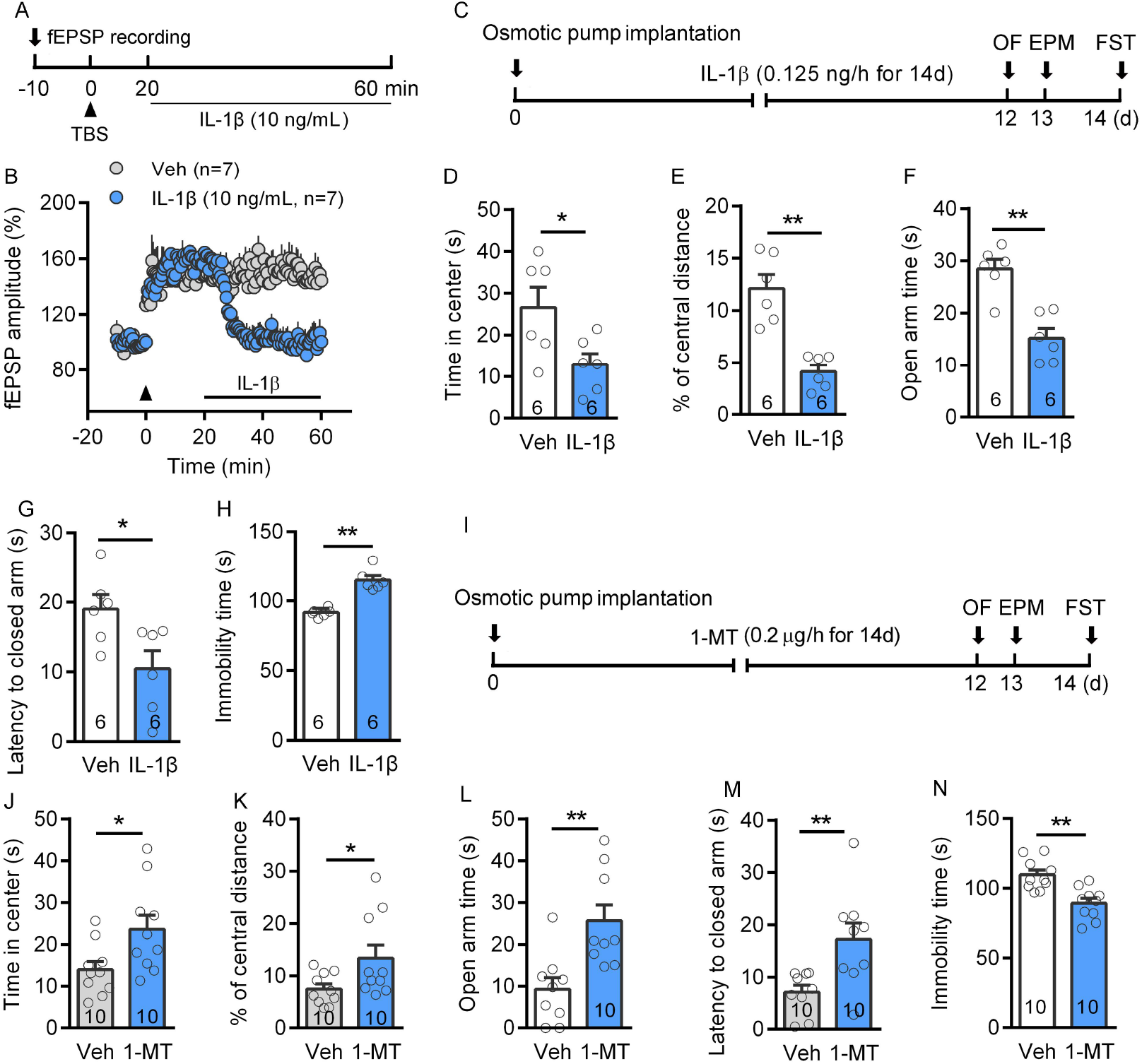
IL-1β and indoleamine 2,3-dioxygenase are involved in CION-induced anxiodepressive-like behaviors. (A and B) Perfusion of IL-1β (5.8 × 10^-10^ mol/L) obviously impairs the hippocampal LTP of naïve mice (n=7/7, vehicle/IL-1β, slices/group). (C) Schematics of the protocol for experiments D-H. (D-H) Intra-CA1 of IL-1β (0.125 ng/h) by osmotic pump system directly results in anxiodepressive-like behaviors in OF (D and E), EPM (F and G) and FS (H) tests in naive rats. \**P*<0.05, \*\**P*<0.01, two-tailed Student’s t test (n=6/6, vehicle/IL-1β, rats/group). (I) Schematics of the protocol for experiments J-N. (J-N) Intra-CA1 of 1-methyl-D-tryptophan (1-MT, 0.2 μg/h), a selective inhibitor of indoleamine 2,3-dioxygenase 1 (IDO1), by osmotic pump system significantly blocked CION-induced anxiodepressive-like behaviors in OF (J and K), EPM (L and M) and FS (N) tests in CION rats. \**P*<0.05, \*\**P*<0.01, two-tailed Student’s t test (n=10/10, vehicle /1-MT, rats/group).

## Discussion

As the primary immune cells in the central nervous system, microglia have been well documented to be involved in the pathogenesis of both depression and chronic pain. The implication of microglia in depression is supported by the magnetic resonance imaging showing the obvious activation of microglia in multiple brain regions of depressed patients (40). Microglia activation in hippocampus was also revealed in multiple animal models of depression induced by chronic stress. For example, Frank el al. reported a priming of microglia in hippocampus of animals exposed to inescapable tailshock (41). Duman el al. demonstrated that chronic unpredictable stress upregulated microglial activity in hippocampal CA3 region and induced depressive-like behaviors (42). A recent study from Engblom’s lab found that microglial activation-induced negative emotion was mediated by prostaglanding-dependent modulation of striatal neurons, suggesting that microglia might be an interference target for treatment of depressive symptoms (10). The causal relationship between microglia activation and neuropathic pain has been well established in the spinal cord level (43, 44), but the role of supraspinal microglia in neuropathic pain is still debated. Spared nerve injury significantly increased the expression levels of IBA-1 and IL-1β on day 12 not only in spinal dorsal horn but also in hippocampus CA1, prefrontal cortex, nucleus accumbens, and amygdala (45). The increased microglial expression in the anterior cingulate cortex (ACC) was also observed at day 7 after sciatic nerve ligation, which can be reversed by intracerebroventricular (i.c.v) treatment with the microglia inhibitor minocycline (46). Similarly, following spinal nerve ligation, hippocampal microglia were robustly activated, and inhibition of hippocampal microglia could relieve neuropathic pain (47). Differently, some studies showed that peripheral nerve injury selectively activated microglia in the spinal dorsal horn cord without affecting the supraspinal structures (48, 49). These contradictions might result from the differences in animal strains or species, animal models and time point of experimental observation etc. Indeed, our current study demonstrated that significant activation of hippocampal microglia occurred only two weeks after CION, not within one week. More importantly, the time window of hippocampal microglial activation coincides with the development of anxiodepressive-like consequences of neuropathic pain, implying the involvement of microglia activation in anxiodepressive consequences of neuropathic pain. Consistent with previous studies (6, 50)), the present results showed that the different symptoms of TN, including mechanical allodynia and anxiodepressive-like consequences, display different time courses following nerve injury. Animals developed mechanical allodynia within 3 days after CION and anxiodepressive-like behaviors occurred after 2 weeks, which allows us to study the time-dependent evolution of hippocampal microglial activation with anxiodepressive-like symptom of TN.

Microglia have been shown to participate in the process of synapse pruning and stripping by selectively contacting synapse with their vicinity, and thus affecting synapse plasticity (51). Evidence has shown that peripheral nerve injury can impair LTP at hippocampal CA3-CA1 synapse by increasing the level of TNF-α in a microglia-dependent mechanism (52, 53). Microglia activation was also proved to direct suppress hippocampal LTP (54). In the present study, we further revealed that TN impaired ipsilateral hippocampal LTP at 2 weeks after CION surgery, when the hippocampal microglia were activated, and anxiodepression-like behaviors were developed. Optogenetic activation of microglia directly impaired hippocampal LTP. On the contrary, optogenetic inhibition of microglia prevented the TN-induced LTP impairment, suggesting a key role of microglia in regulating hippocampal synapse plasticity.

P2X7R is an ATP-gated, nonselective cation channel that widely expressed in immune-associated cells, including macrophages, mast cells and microglia (55). Previous studies from our and other laboratories showed that in the spinal cord P2X7R was preferentially expressed in microglia and play a pivotal role in the cross talk between microglia and neurons (36, 56, 57). Herein, we further demonstrated P2X7R upregulation in the hippocampal microglia in chronic pain with anxiodepressive status. Knockdown of P2X7R inhibited CION-induced IBA-1 upregulation and anxiodepressive-like behaviors. Analogously, P2X7R antagonist BBG also reversed the microglial activation in cortical, hippocampal regions and the basal nuclei of mouse brains in an unpredictable chronic mild stress (UCMS) model (37, 58). In the absence of pathological insults, overexpression of P2X7R is sufficient to drive the activation and proliferation of microglia in rat primary hippocampal cultures. P2X7R specific antagonist, oxidized ATP (oxATP), was effective in markedly attenuating microgliosis (59). As the resident immune cells of central nerve system, microglia could initiate a pro-inflammatory response by detecting the upregulation of damage-associated molecular patterns, such as ATP, in the microenvironment (60, 61). In recent years, activated microglia has been demonstrated to participate in the pathogenesis of multiple neuroinflammation-related diseases including depressive disorder by secreting IL-1β and other inflammatory cytokines (62–64). Activation of P2X7R on murine and human neutrophils exacerbated inflammatory responses by NLRP3 inflammasome-dependent IL-1β secretion (65). Our previous study has revealed that P2X7R-mediated NLRP3 inflammasome assembly in hippocampal microglia contributes to UCMS-induced depression (37). In the central nervous system, the key player involved in the secretion of biologically active IL-1β may is P2X7R (66). The current study showed that CION-induced IL-1β upregulation in hippocampal CA1 area was blocked by p*2x7r*-siRNA; Microglial activation-induced anxiodepressive-like behaviors and hippocampal LTP impairment were prevented by IL-1Ra. Taken together, it could indeed be authentic that P2X7R mediated hippocampal microglial purinergic inflammatory responses leading to IL-1β increase, which may be a pathogenesis of anxiodepression caused by TN (Supplemental Figure 8).

The roles of hippocampus in the pathogenesis of depression has been well established. However, most experimental observations and functional manipulation are directed at the bilateral hippocampus. Unconventionally, we found that long-term CION exposure resulted in microglial activation in the ipsilateral but not contralateral hippocampus. CION-induced LTP impairment occurred only on the ipsilateral hippocampus. Pharmacological or optogenetic inhibition of ipsilateral hippocampal microglia significantly blocked trigeminal neuralgia-induced anxiodepressive-like behaviors. Optogenetic activation of unilateral hippocampal microglia is sufficient to evoke anxieodepressive-like states. These results suggest that the role of bilateral hippocampal CA1 area in trigeminal neuralgia-induced anxiodepression is asymmetric. As early as 1994, it had been reported that activation of the 5-HT1Areceptor in the right hippocampus rather than left hippocampus produced anxiety (67). Delivering vasoactive intestinal peptide (VIP) into left or bilateral hippocampus exhibited obvious antinociceptive effects in olfactory bulbectomy rats, a widely used animal model of depression, whereas delivering VIP into right hippocampus has no effects (68). Recently, Hodaie’s laboratory found that right-side trigeminal neuralgia patients have significant volumetric reductions in ipsilateral CA1, CA4, dentate gyrus and hippocampus-amygdala transition area, compared with healthy controls (69). Our recent study in proteomic analysis found that differences in the biomolecules and signaling pathways between ipsilateral and contralateral hippocampus 14 days after unilateral CION were evident. WASL protein that may play an important role in microglia activation (70) was upregulated only in the ipsilateral hippocampal CA1 area (71). In addition, trigeminal ganglion (TG) sensory neurons have been revealed to provide a direct monosynaptic input to ipsilateral lateral parabrachial nucleus (PB_L_), and activating this ipsilateral TG→PB_L_ monosynaptic projection in awake animals can elicit robust aversive behaviors (72). As a key relay node in the affective pain circuit, PB_L_ relays the nociceptive afferent information to multiple limbic regions, including the central amygdala, the anterior cingulate, the lateral hypothalamus and the hippocampus (73–75). Thus, the ipsilateral projection of TG neurons to brain provides an anatomical basis for the connection between facial nociceptive inputs and emotion-related brain regions, which may partly explain why ipsilateral hippocampus play an important role in the trigeminal neuralgia-induced anxiodepression.

In summary, using genetic and pharmacological bidirectional manipulation of microglial activity in the hippocampal CA1 area, we found that ipsilateral activation of microglia is necessary for TN-induced anxiety and depression; and unilateral activation of hippocampal microglia is sufficient to induce an anxiodepressive state in rodents. Microglial purinergic inflammatory responses and the neuronal plasticity changes in the ipsilateral hippocampal CA1 area may be a key pathogenesis of anxiodepression caused by TN. The approaches targeting microglia and P2X7R signaling might offer novel therapies for chronic pain-related anxiety and depressive disorder.

## Methods

### Experimental animals

Adult Wistar rats (190-220 g) and C57BL/6 (10-12 weeks) were purchased from Shanghai Experimental Animal Center of Chinese Academy of Science. P2X7R KO mice (JAX#005576), CX3CR1-CreER-GFP (JAX#021160), ChR2-Flox (JAX#012735) and Arch-Flox (JAX#012567) mice were used in this study (male Wistar and C57BL/6J, both male and female transgenic strains). CX3CR1-GFP::ChR2 and CX3CR1-GFP::Arch mice were bred by CX3CR1-CreER-GFP and ChR2-Flox and Arch-Flox mice respectively. All animals were housed in a temperature and humidity controlled room on a 12h/12h light-dark cycle and access to food and water ad libitum. The body weight and sexes of transgenic animals were assigned to different treatment groups randomly. All the following behavioral testing and electrophysiological recording and quantification of immunohistochemistry and western blot experiments described herein were performed by experiments who were blind to the treatments.

### Trigeminal neuralgia surgery

Trigeminal neuralgia (TN) model was established by chronic constriction injury of the unilateral infraorbital nerve (CION) via an intraoral approach as described previously (76). In brief, rats or mice were anesthetized with sodium pentobarbital (50 mg/kg, i.p.), and the head was fixed, keeping the body supine and mouth wide open. A surgical incision was made at 0.1 cm proximal to the first molar along the left gingivobuccal margin, and the left or right infraorbital nerve was exposed. Two (for rat) or one (for mouse) ligatures with 4-0 chromic gut ligatures were tied loosely around the nerve at approximately 1.5 mm apart. Sham-operated animals received only nerve exposure but not ligation. Subsequently, the overlaying mucosa was closed. All surgical procedures were performed aseptically.

### Intra-CA1 drug infusions

Rats were anesthetized with intraperitoneal (i.p.) injection of sodium pentobarbital (50 mg/kg), and then securely placed into a rat stereotaxic device with bregama and lambda horizontally level. A stainless steel cannula (OD 0.41 × ID 0.25, RWD Life Science Co., Ltd, Shenzhen, China) with a stainless steel stylet plug (OD 0.21 × ID 0.11, RWD Lifer Science) was unilaterally or bilaterally implanted 0.5 mm above the CA1 of dorsal hippocampus injection site [from bregma: anteroposterior (AP) -3.2 mm, mediolateral (ML) ±1.8 mm; dorsoventral (DV) -2.5 mm] according to the rat brain’s altas (77). The cannula was fixed to the cranium with denture acrylic cement. Animals were allowed to recover for 7 days. Microinjection was carried out under isoflurance brief anaesthesia through a 33 gauge stainless steel injection cannula that extended 0.5 mm beyond the tip of the guide cannula. The injection cannula was connected to a 1-μL Hamilton syringe through PE-10 tubing. A volume of 0.6 μL of either vehicle or drug was injected with a steady speed (0.1μl /min). Small interfering RNA (siRNA) targeting the rat P2X7R mRNA (L-091415-00-0020) or nontargeting control siRNA (D-001220-01-20) was mixed with polyethylenimine (PEI; 1.8 μL of 10 mM PEI/μg RNA), as we previously demonstrated (36). Mac-1-saporin-toxin (15 μg in 8.8 μL) and saporin control (8.8 μg in 8.8 μL) were purchased in solution from Advanced Targeting Systems. Doses were determined from pilot experiments. The injection cannula was left in the place for additional 5 min to minimize drug overflow along the injection track. At the end of the experiment, brains were sectioned for neutral red staining to verify cannula position and injection site.

### Implantation of micro-osmotic pump

Micro-osmotic pumps were used for continuous hippocampal CA1 delivery. Implantation of the unilateral or bilateral hippocampal CA1 cannula was performed as described above. A micro-osmotic pump (Model 1002, Alzet, Cupertino, CA, USA) with a delivery rate of 0.25 μL/h during 14 days, connected via polyethylene tubing to a brain infusion kit (Alzet), was embedded under the skin of the neck. A740003, IL-Ra, IL-1β and 1-MT were delivered in a dosage of 9 μg/d, 5μg/d, 3 ng/d and 5μg/d during 14 days, respectively. 1-MT, IL-Ra and IL-1β were dissolved in 0.9 % saline, and A740003 were dissolved in 10 % dimethyl sulfoxide (DMSO).

### Optogenetics experiments

CX3CR1-GFP::ChR2 and CX3CR1-GFP::Arch mice were used in the optogenetic experiments. Optical fibers (Newdoon Technology Co., Ltd, Hangzhou, China) were unilaterally implanted upon the mouse hippocampal CA1 (from bregma: AP, -2.05 mm; ML, ±1.25 mm; DV, -1.05 mm). Experiments were performed at one week after optical fiber implantation. Tamoxifen (50 mg/kg/day, i.p.) was injected into CX3CR1-GFP::Arch mice before CION, or CX3CR1-GFP::ChR2 mice for three days. For the experiments of microglia inhibition, yellow light (580 nm, 6 mW, 50 s/on, 10 s/off, continuously, 1h/day, for 8 days) was delivered into the ipsilateral hippocampus from day 8 to day 15 after CION. For the experiments of microglia activation, blue light (473 nm, 6 mW, pulse width 5 ms, 20 Hz, 30 min/day) was delivered into the unilateral hippocampus for three continuous days.

### *von* Frey test

Each animal was handled and habituated to the testing environment for 30 minutes at least 3 days before testing. Rats were placed into a customized cage individually, and a series of calibrated *von* Frey filaments (2, 4, 6, 8, 10, 15, 26 g, Stoelting Company, Wood Dale, Illinois, USA) were lightly applied to the skin within the infraorbital territory, near the center of the vibrissal pad on hairy skin surrounding the mystacial vibrissae. The mouse was gently held by an experimenter wearing a regular leather work-glove, and mechanical sensitivity was determined with a series of *von* Frey hairs (0.07, 0.16, 0.4, 0.6, 1, 1.4, 2, 4 g). The filaments were performed in an increasing order from the lowest force, and a brisk or active withdrawal of the head from the probing filament was defined as a response. Each filament was tested 5 times at 5-s intervals. The withdrawal threshold was defined as the lowest force in grams that produced at least 3 withdrawal responses in 5 consecutive applications.

### Assessment of cold allodynia

After acclimatized to the experimental environment for 30 min, the rat was gently grasped and restrained by tester with leather work-glove for at least 10 min to minimize stress. Acetone (90%, 30 μL) was dropped onto the unilateral vibrissal pad skin, using a 25-gauge blunt needle attached to a 50 μL micro-syringe. Immediately after acetone administration, the rat was placed back to their home cage and the nociceptive behavior was observed within 5 min. Asymmetric orofacial grooming was defined as nociceptive behaviors, and the time animals spent in this behavior was recorded.

### Open field test (OFT)

The apparatus consisted of an open box (length × width × height: 60 cm × 60 cm × 35 cm for rat; 40 cm × 40 cm × 30 cm for mouse). The observation arena was divided into the angle zone (15 cm × 15 cm at the 4 corners for rat; 10 cm × 10 cm at the 4 corners for mouse), the center of the arena (30 cm × 30 cm for rat; 20 cm × 20 cm for mouse). The house was maintained in a dim illumination (25 lux) with no noise. Rats or mice were gently placed into the center of the arena and allowed to explore for 5 minutes. Video tracking software (Ethovision XT v11.5, Noldus BV) was used to record the distance traveled in each zone by the animal.

### Elevated plus maze (EPM)

The EPM consisted of 4 arms (10 cm × 50 cm for rats; 6 cm × 35 cm for mice) and a central platform (10 cm × 10 cm for rats; 6 cm × 6 cm for mice) elevated 50 cm above the floor. Two closed arms were enclosed the 45 cm-high (rats) or 10 cm-high (mice) walls crossing with 2 open arms (without walls). The maze was placed in a room with an illumination of 25 lux. Animals were placed in the central of the maze facing an open arm and were allowed to freely explore the maze for 5 minutes. Both the open/closed arm time and latency of the first entry into the closed arm were recorded by the video tracking system (Ethovision XT v11.5, Noldus BV).

### Forced swimming test (FS)

Animals were placed into a clear cylindrical glass container (high × diameter, rats: 60 cm × 30 cm; mice: 45 cm × 20 cm) filled with water (24℃± 2℃) to a depth adjusted for the weight of the individual animal, so that its hind paws could just touch the bottom of the container. Animals were trained for 10 minutes on the first day, then animals were replaced in the cylinders on the second day. The total duration of immobility and activities of the animals were recorded during a 5-min test by the video. Immobility was defined as the animal not making any active movements other than that necessary to keep the head and nose above the water. The animals were dried immediately and returned to their home cages after the test.

### Rotarod test

A rotarod test was used to determine whether CION surgery influences the motor function of animals. Animals were placed on an accelerating rotating rod (IITC Life Science), and the rotating rod underwent a liner acceleration from 0 to 40 rounds per minute (rpm) within 5 min. Animals were trained for three trails with a break of 10 min between the trails per day for 2 consecutive days before test. On day 3, animal behaviors on the accelerating rotating rod was scored for their latency (in seconds) to fall for each trial.

### Western blotting

Animals were sacrificed with overdoses of urethane, and the hippocampal CA1 area were rapidly isolated. The hippocampus tissues were homogenized in lysis buffer (12.5 μL/mg) containing a mixture of protease inhibitors and phenylmethylsulfonyl fluoride (Roche Diagnostics, Indianapolis, IN). The protein concentration was determined by a BCA protein assay kit (Pierce, Rockford, IL) according to its instruction provided by manufacturers. Protein samples (∼15 μg) were loaded and separated on 10% sodium dodecyl sulfate polyacrylamide gel electrophoresis (SDS-PAGE, Bio-Rad, Hercules, CA), and transferred to polyvinilidene difluoride membranes (PVDF, Millipore, Billerica, MA). After blocked with 10% non-fat milk at room temperature (RT) for 2 h, the membranes were incubated overnight at 4℃ with primary antibodies, followed by horseradish peroxidase (HRP)-conjugated secondary antibodies (1:10000; Pierce, Rockford, IL) for 2 h at 4 ℃. GAPDH antibody was probed as a loading control. Signals were detected by enhanced chemiluminescence (Pierce) and captured by ChemiDox XRS system (Bio-Rad). We used the following primary antibodies: rabbit anti-P2X7R (1:2000, #APR-004, Alomone labs, Jerusalem, Israel), goat-anti IBA-1 (1:1000, #ab5076, Abcam, Cambridge, MA), and rabbit-anti IL-1β (1:500, rabbit, PeproTech, Rocky Hill, NJ). All the western blot analysis was performed three to four times, and consistent results were obtained. A Bio-Rad image analysis system was then used to measure the integrated optic density of the specific bands.

### Immunohistochemistry

Animals were deeply anesthetized with overdoses urethane and were transcardially perfused with normal saline followed by pre-cold 4% paraformaldehyde in 0.1 M phosphate buffer (PB, pH 7.4). The brain was removed carefully and postfixed in 4% paraformaldehyde for additional 8-12 h at 4℃. After dehydrated with gradient (10-30%) sucrose in PB at 4℃, Coronary sections (30 μm) were cut on a cryostat microtome (Leica CM 1950, Germany). The sections containing hippocampus were blocked with 10% donkey serum with 0.3% Triton X-100 for 2 h at RT and incubation overnight at 4℃ with corresponding primary antibodies: goat anti-IBA-I (1:1000, Abcam, Cambridge, UK), chicken anti-YFP/GFP (1:500, Aves, Tigard, Oregon), rabbit anti-CD68 (1:500, Abcam) or rabbit anti-P2X7R (1:500, Alomone labs, Israel). The sections were then incubated for 2 h at 4℃ with Alexa fluor 488-conjugated secondary antibody (1:200, Invitrogen, #A21206, Carlsbad, CA, USA) or DAPI (1:30000, Sigma, #32670). For double immunofluorescence, the sections were incubated with a mixture of rabbit anti-P2X7R (1:500) and goat anti-IBA-I (1:1000)/mouse anti-GFAP (1:2000, #G6171, Sigma-Aldrich, St. Louis, MO)/rabbit anti-NeuN (1:2000, #MABN140, Millipore, Billerica, MA) primary antibodies. The sections were then incubated with a mixture of Alexa fluor 488- and 546-conjugated secondary antibodies (1:200) for 2 h at 4℃. The specificity of immunostaining and primary antibodies was verified by omitting the primary antibodies and gene knock-out mice (e.g., P2X7R knock-out mice). The stained sections were observed and images captured with a confocal laser-scanning microscope (Model FV1000, Olympus, Japan).

### Microdialysis and ATP assay in the hippocampus

The microdialysis probe (MD-2211, BASi, West Lafayette, IN) was inserted into the bilateral hippocampal CA1 area via the guide cannula (MD-2255, BASi) to 1 mm beyond the tip of the guide cannula. The dialysis probe was connected to a microinfusing pump (BAS, Microdialysis Syringe 1.0 mL). The probe was perfused with artificial cerebrospinal fluid (ACSF) at a flow rate of 2 μL/min. After dialysate levels stabilized (about 1 h), sample was collected. To prevent degradation of ATP, ecto-ATPase (1 mmol/20 μL; Sigma-Aldrich) was applied as the perfusate. The ATP concentration was measured using an ENLITEN ATP Assay System with a bioluminescence detection kit (Promega, Madison, WI). Briefly, samples (90 μ L) were neutralized to pH 7.4 with 10 μL of 4 M Tris. The luciferase reagent was added 1 s before a 5 s measurement in a luminometer, as described by the supplier (Promega). Light photons were measured by the luminometer and compared with the standard curve to calculate ATP concentration.

### Hippocampal slice preparation

Coronal brain slices containing the hippocampus were obtained from mouse. After anesthetizing with isoflurane, mice were decapitated. The brain was quickly removed and immediately submerged in pre-oxygenated (95 % O2, 5 % CO2) cold cutting solution (93 mM N-methyl-D-glucamine, 93 mM HCL, 2.5 mM KCL, 1.2 mM NaH2PO4, 30 mM NaHCO3, 20 mM HEPES, 25 mM glucose, 5 mM sodium ascorbate, 2 mM Thiourea, 3 mM sodium pyruvate, 10 mM MgSO4.7H2O, 0.5 mM CaCl2.2H2O, and 12 mM N-Acetyl-L-cysteine). The osmolarity was adjusted to 290-320 mOsmol/L and the pH to 7.3-7.4. A tissue block containing the hippocampus was fixed on the stage using cyanoacrylate glue, and then cut into 350 μm sections by a vibratome (Leica VT 1200S, Leica, Germany). Subsequently, hippocampal slices were transferred to an oxygenated chamber filled with holding ACSF (94 mM NaCl, 2.5 mM KCl, 1.2 mM NaH2PO4, 30 mM NaHCO3, 20 mM HEPES, 25 mM glucose, 5 mM sodium ascorbate, 2 mM Thiourea, 3 mM sodium pyruvate, 2 mM MgSO4.7H2O, 2 mM CaCl2.2H2O, and 12 mM N-Acetyl-L-cysteine) at RT for 30 min, and then transferred to regular ACSF (126 mM NaCl, 4.0 mM KCl, 1.25 mM MgCl2, 26 mM NaHCO3, 1.25 mM NaH2PO4, 2.5 mM CaCl2, and 10 mM glucose) at RT for 1 h before recording.

### Electrophysiological recording

A single hippocampal slice was transferred to a recording chamber and continuously perfused with regular ACSF at a rate of 5 mL/min at RT. For recording field excitatory postsynaptic potentials (fEPSPs), a single glass pipette filled with 4 M NaCl was used and a bipolar tungsten stimulating electrode placed in CA1 stratum radiatum of hippocampus was used to stimulate excitatory response at Schaffer collateral -CA1 synapses. Electrical stimuli (100 μs pulses) were delivered to Schaffer collateral fibers by a square pulse stimulator (Master-8; AMPI, Jerusalem, Israel) and a stimulus isolator (ISO-Flex; AMPI) at an interval of 30 s. Stimulation intensities were adjusted to evoke approximately half of the maximal field potentials. The specific fEPSP signals were acquired using a patch clamp amplifier (Axopatch 700B; Axon Instruments, Foster City, CA, USA) and digitized with an axon Digidata 1440A. After obtaining a stable baseline of 10 min, long term potential (LTP) was induced by a theta burst stimulation (TBS, 4 bursts of 4 pluses at 100 Hz delivered at 200 ms intervals, repeated 4 times with intervals of 10 s). Following TBS, the intensity was then adjusted to the level previously utilized to produce baseline fEPSPs and recorded for additional 1 h.

### ELISA assay

Hippocampal slices were prepared as described above under sterile condition, were maintained in sterile culture medium at cell incubator with 95% O2 and 5% CO2 for two weeks. Hippocampal slices were treated with different agents: LPS (100 ng/ml), BzATP (μmol/L) + LPS (100 ng/ml) and A740003 (100 μmol/L) + BzATP (μmol/L) + LPS (100 ng/ml). Twenty-four hours later, hippocampal slices were collected and homogenized in the lysis buffer (12.5 μL/mg) containing protease and phosphatase inhibitors (Sigma Chemical Co) at 4℃. Then, the tissue lysate was centrifuged with 12000 rpm for 15 min to get the supernatant. ELISA was conducted according to manufacturer’s instructions (R&D Systems, #RLB00, Minneapolis, MN) to assay IL-1β content in the supernatant.

### QUANTIFICATION AND STATISTICAL ANALYSES

#### Microglia morphological analysis

Hippocampal microglia were stained with IBA-1 and imaged with a confocal laser-scanning microscope at 60× magnification. The Neurolucida software (MBF Bioscience, Williston, VT) was applied for three-dimensional (3D) reconstruction of microglia within CA1 area of hippocampus. Sholl analysis was performed to analyze the morphology of microglia by placing 3D concentric circles in 5-mm increments starting at 5 mm from the soma using the NeuroExplorer software.

#### Statistical analysis

All data were presented as the mean ± SEM. No statistical power calculation was conducted before the study. The sample sizes were based on our previous knowledge and experience with this design. Animals that showed hyperactivity and lethargy in behavioral tests were excluded from the experiments. All data from different groups were verified for normality and homogeneity of variance using Kolmogorov-Smirnov and Brown-Forsythe tests befor analysis. Behavioral data, electrophysiological recording, immunohistochemistry and western blot data were analyzed using 2-tailed Student’s *t* test when comparing two groups, or 1-way ANOVA followed by post hoc Dunnett’s test or 2-way repeated-measures (RM) ANOVA followed by post hoc Bonferroni’s multiple-comparisons test when comparing more than two groups. No data were excluded from statistical analysis due to outlier status. All the hypothesis testing was two-tailed with *P* value less than 0.05 considered statistically significant. The statistical analysis was performed using GraphPad 7.0 software.

#### Study approval

All the animal experimental procedures were approved by the Committee on the Use of Animal Experiments of Fudan University (permit No. SYXK 2009-0082) and in accordance with policies on the use of laboratory animals issued by the International Association for the Study of Pain (IASP, Washington, D.C.).

## Author contributions

Y.Q.Z. conceived the project and supervised all experiments. L.Q.C., X.J.L. and Q.H.G. performed behavioral experiments. L.Q.C. and S.S.L. performed electrophysiological recording. L.Q.C. and X.J.L. performed immunohistochemistry and optogenetic experiments. L.Q.C. and Q.H.G performed ELISA experiment. Q.H.G. performed microdialysis and western blotting experiments. L.Q.C., J.Y., W.D.X. and Y.Q.Z. analyzed the data. L.Q.C. and Y.Q.Z. wrote the manuscript. All the authors read and discussed the manuscript.

## Acknowledgements

We thank Dr. Lan Ma (Fudan University) for giving some Arch-Flox mice and Dr. Jiayi Zhang (Fudan University) for giving ChR2-Flox mice. This work was supported by grants from National Natural Science Foundation of China (31930042, 82130032, 82021002) and by funds from the innovative research team of high-level local universities in Shanghai, Shanghai Municipal Science and Technology Major Project (No.2018SHZDZX01), ZJLab and Shanghai Center for Brain Science and Brain-Inspired Technology.

