## Supplemental information for "Asymmetric activation of microglia in the hippocampus drives anxiodepressive consequences of trigeminal neuralgia"

### Supplemental Figures

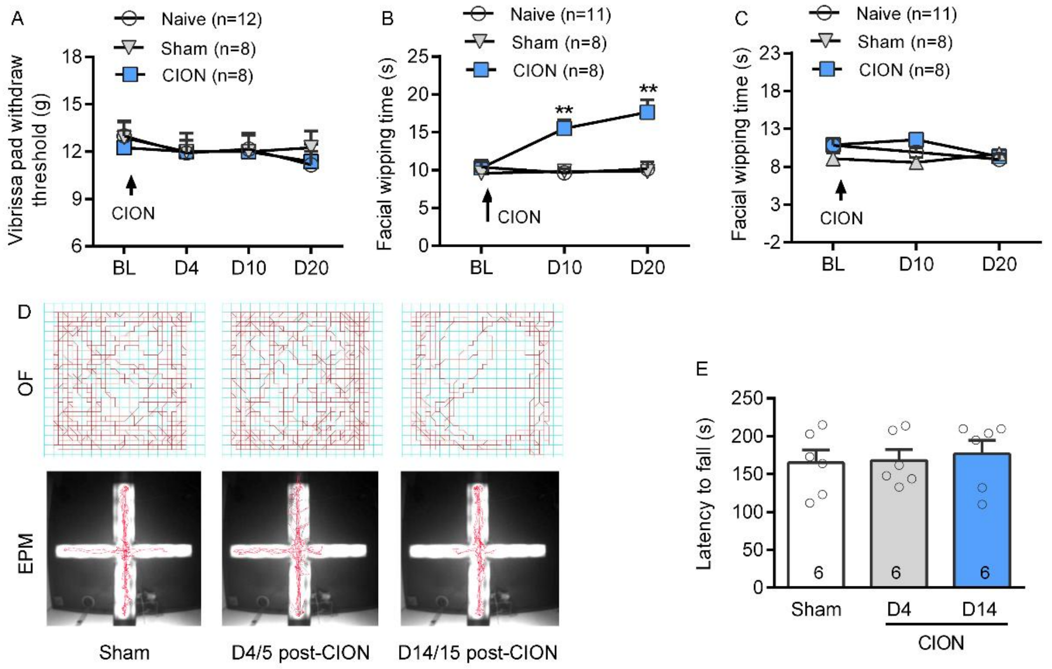


**Supplemental Figure S1. related to Figure 1, Trigeminal neuralgia causes cold allodynia and anxiety-like behaviors.** (A) The mechanical response threshold of the contralateral vibrissa pad is not affected by unilateral TN, remaining stable for 20 days after CION. Two-way RM ANOVA (n=12/8/8, naive/sham/ CION, rats/group). (B and C) Following TN, the duration of facial wiping caused by acetone is significantly prolonged in the ipsilateral (B), but not contralateral vibrissa pad (C). ***P* < 0.01 versus sham and naive, 2-way RM ANOVA followed by post hoc Student-Newman-Keuls test (n=11/8/8, naive/sham/CION, rats/group). (D) Example track plots from sham and CION rats in OF (above) and EPM (below) tests. (E) TN do not influence motor coordination in sham, CION-4day and CION-14 day rats in a rotarod test. One-way ANOVA (n=6 rats for all the groups).

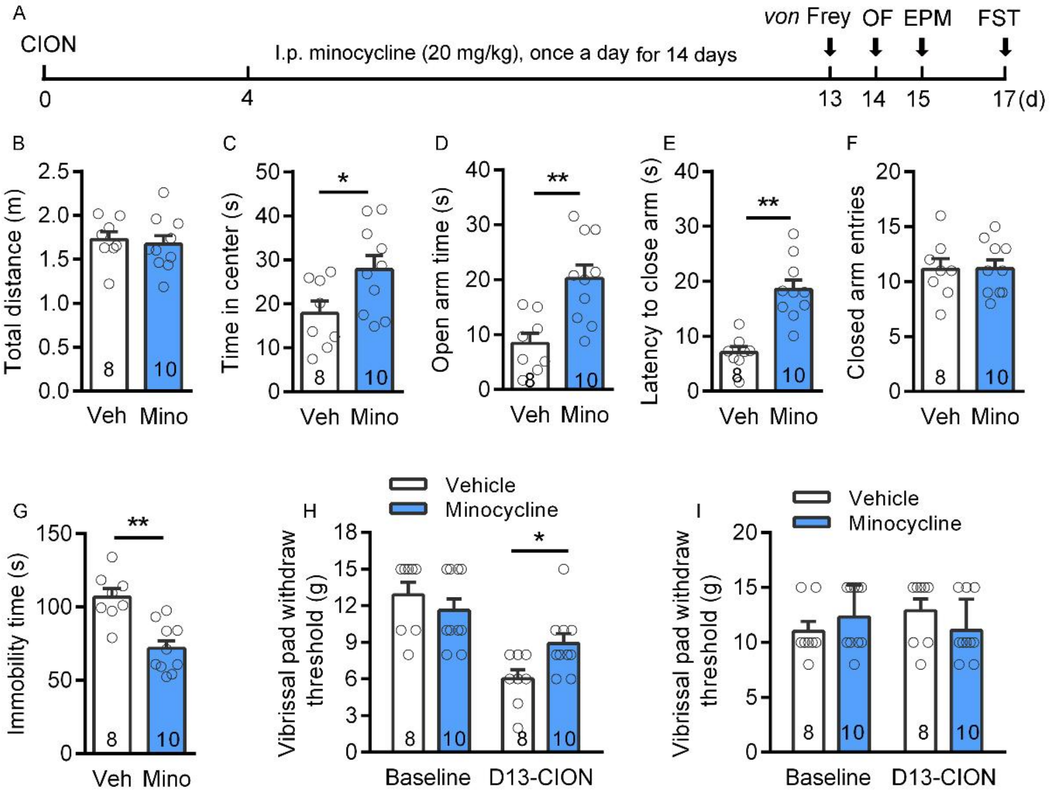


**Supplemental Figure 2. related to Figure 3 and 4, Systemic application of minocycline prevented the development of anxiodepressive-like behaviors and mechanical allodynia caused by CION in rats.** (A) Schematic of the protocol for the experiments B-I. (B-G) Systemic administration of microglial inhibitor minocycline (20 mg/kg once daily for 14 days) completely prevents the CION-induced anxiodepressive-like behaviors in OF (B and C), EPM (D-F) and FS (G) tests. **P*<0.05, ***P* < 0.01, two-tailed Student's *t* test (n=8/10, vehicle/minocycline, rats/group). (H and I) Systemic administration of minocycline partially blocks CION-induced mechanical allodynia in ipsilateral vibrissa pad (H), but does not affect the mechanical response threshold in contralateral vibrissa pad (I). **P*<0.05, two-tailed Student's *t* test (n=8/10, vehicle/minocycline, rats/group).

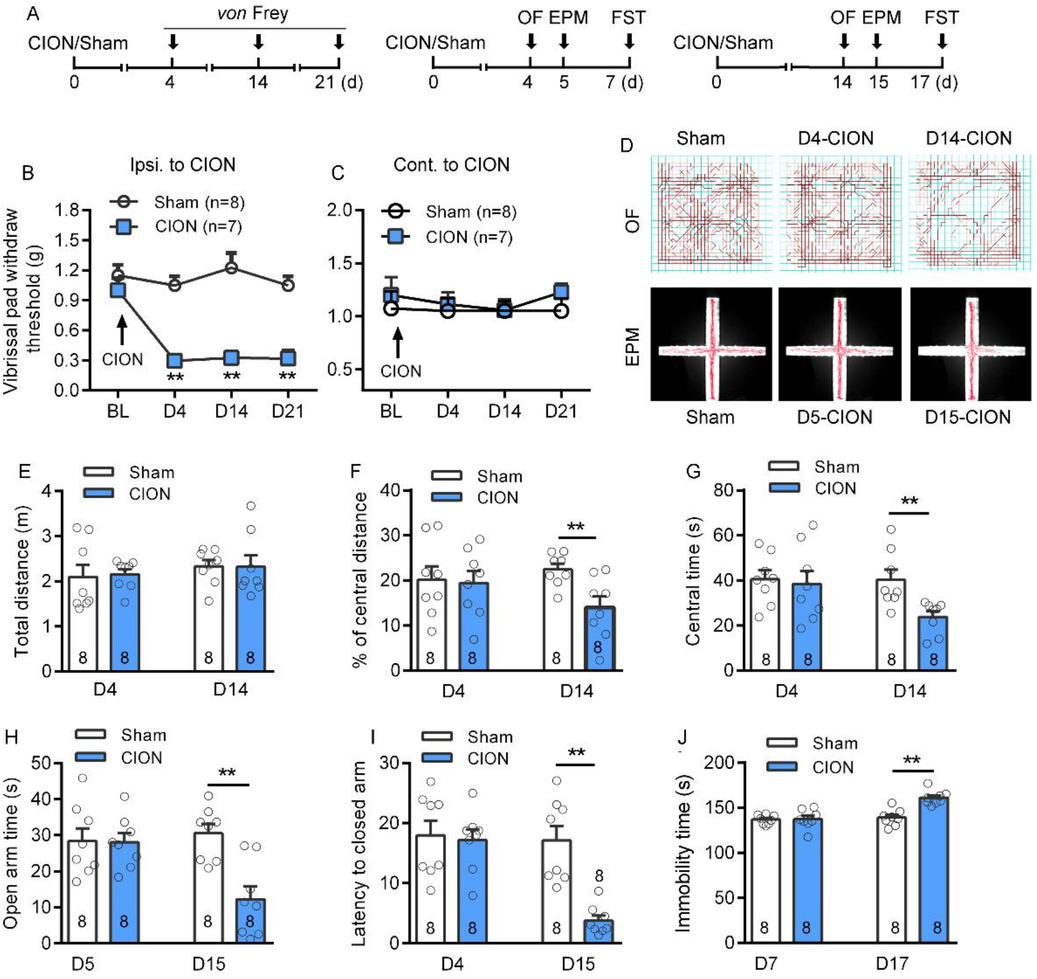


**Supplemental Figure 3. related to Figure 4 and 5, Trigeminal neuralgia time-dependently induces mechanical allodynia of the ipsilateral vibrissa pad and anxiodepressive-like behaviors in mice.** (A) Schematic of the protocol for the experiments B-J. (B and C) Following the trigeminal neuralgia (TN), mechanical response threshold of the ipsilateral, but not contralateral vibrissa pad, significantly decreases on day 4 and lasted for more than 21 days in C57 mice. $^{**}P < 0.01$ versus sham, 2-way RM ANOVA followed by post hoc Student-Newman-Keuls test (n=8/7, sham/CION, mice/grou). (D) Example track plots from sham and CION mice in OF (above) and EPM (below) tests. (E-J) The TN mice exhibits anxiodepressive-like behaviors in OF (E-G), EPM (H and I) and FS (J) tests on days 14-17 but not days 4-7 after CION. $^{**}P < 0.01$, 1-way RM ANOVA followed by post hoc Student-Newman-Keuls test (n=8 mice for all the groups).

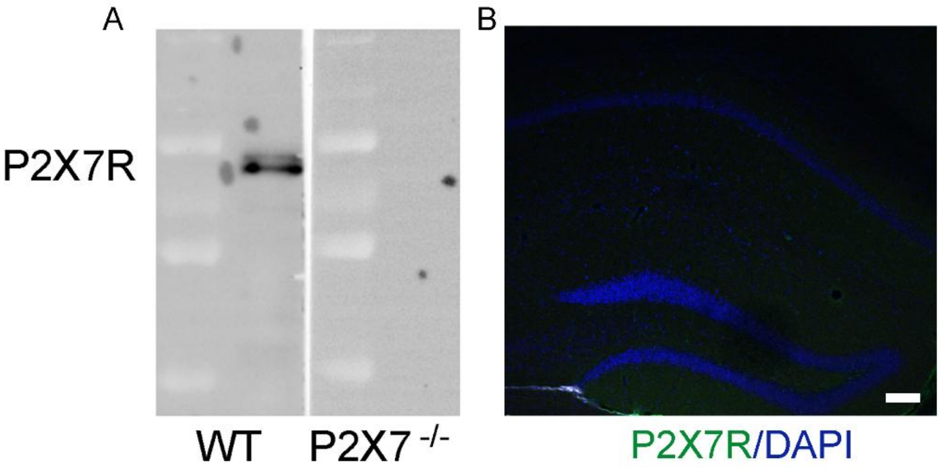


**Supplemental Figure 4. related to Figure 6, Verification of the specificity of P2X7R antibody.** (A) Western blot of hippocampal tissue lysates from WT and P2X7 KO mice using anti-P2X7R antibody. (B) Immunofluorescence staining reveals no P2X7R-IR is detected in the hippocampus of P2X7 KO mice. Scale bar indicates 100 µm.

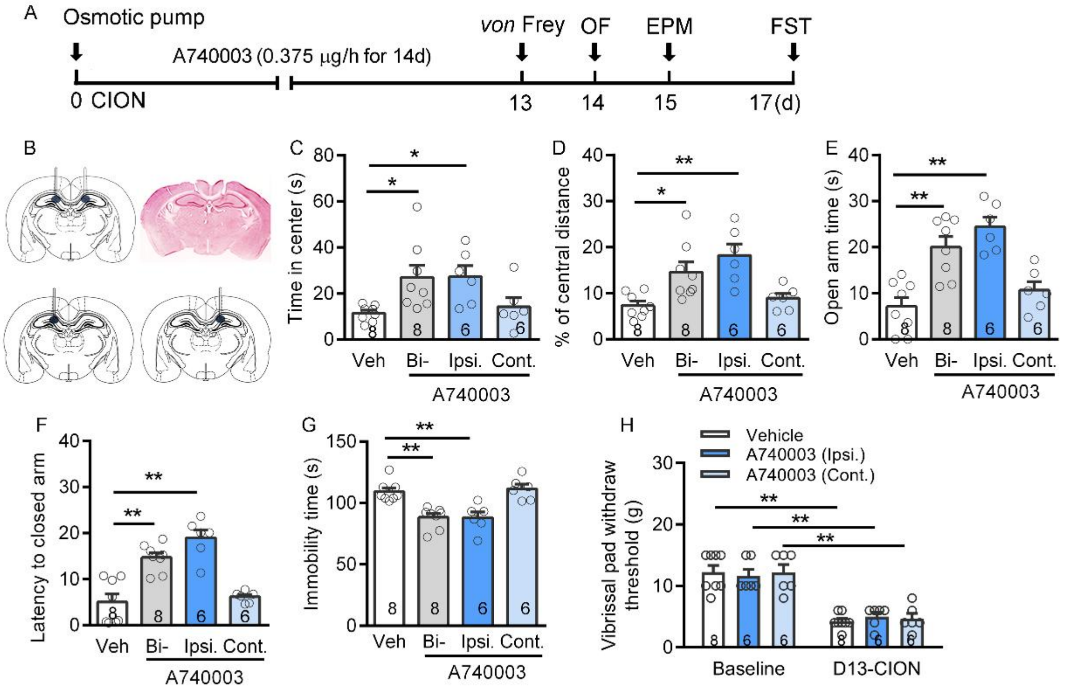


**Supplemental Figure 5. related to Figure 6, Blockade of P2X7R in the bilateral or ipsilateral hippocampus reverses the CION-induced anxiodepressive-like behaviors.** (A) Schematic of the protocol for the experiments C-H. (B) Schematic and photomicrograph of coronal section showing cannula placement in the bilateral and unilateral hippocampus. (C-G) Ipsilateral or bilateral injections of A740003 (a P2X7R specific antagonist, 9 μg/d) into the CA1 area of hippocampus by osmotic pump system lead to a significant anti-anxiodepressive effect in OF (C, D), EPM (E, F) and FS (G) tests in CION rats, whereas administration of A740003 into the contralateral hippocampal CA1 fails to block the development of anxiodepressive-like behaviors. $^*P<0.05$, $^{**}P<0.01$, 1-way ANOVA followed by post hoc Student-Newman-Keuls test (n=8/8/6/6, vehicle/ bilateral A740003/ipsilateral/contralateral A740003, rats/group). (H) Either ipsilateral or contralateral intra-CA1 of A740003 does not affect CION-induced mechanical allodynia. $^{**}P<0.01$, 1-way ANOVA followed by post hoc Student-Newman-Keuls test (n=8/6/6, vehicle/ipsilateral/contralateral A740003, rats/group).

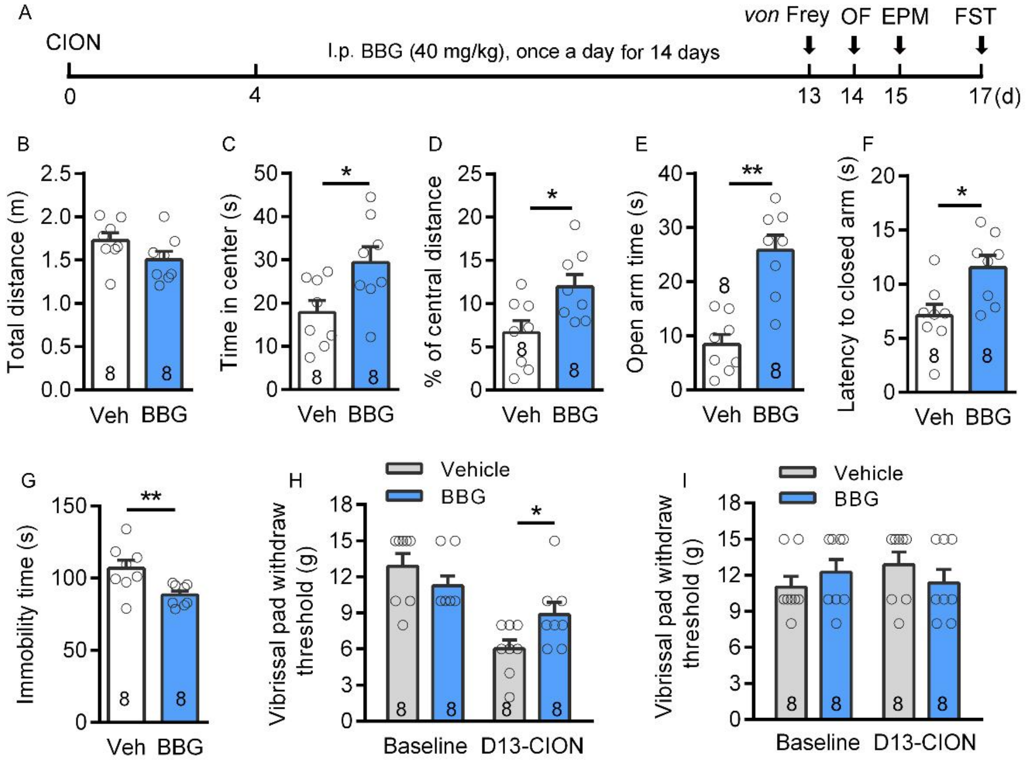


**Supplemental Figure 6. related to Figure 6, Systemic application of brilliant blue G (BBG) prevented the development of anxiodepressive-like behaviors and mechanical allodynia caused by CION in rats.** (A) Schematic of the protocol for the experiments B-I. (B-G) Systemic administration of BBG significantly blocks the CION-induced anxiodepressive-like behaviors in OF (B-D), EPM (E and F) and FS (G) tests. **P*<0.05, ***P* < 0.01, two-tailed Student's *t* test (n=8/8, vehicle/BBG, rats/group). (H and I) Systemic administration of BBG (40 mg/kg once daily for 14 days) partially blocks CION-induced mechanical allodynia in ipsilateral vibrissa pad (H), but does not affect the mechanical response threshold in contralateral vibrissa pad (I). **P*<0.05, 1-way ANOVA followed by post hoc Student-Newman-Keuls test (n=8/8, vehicle/BBG, rats/group).

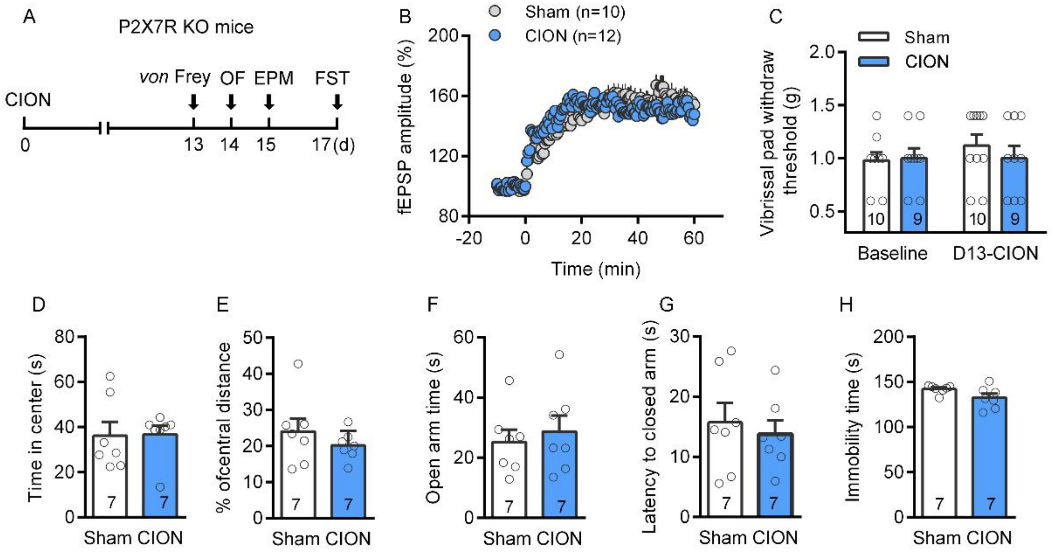


**Supplemental Figure 7. related to Figure 6, P2X7R knockout (P2X7R KO) mice are resistant to CION-induced mechanical allodynia and anxiodepressie-like behaviors.** (A) Schematic of the protocol for the experiments B-H. (B) Trigeminal neuralgia does not impair hippocampal LTP on day 14 after CION in P2X7R KO mice (n=10/12 sham/CION; slices/group). (C) Trigeminal neuralgia do not induce mechanical allodynia of vibrissa pad. Two-tailed Student's *t* test (n=10/9, sham/CION, mice/group). (D-H) Trigeminal neuralgia fails to induce anxiodepressie-like behaviors. Two-tailed Student's *t* test (n=7/7, sham/CION).

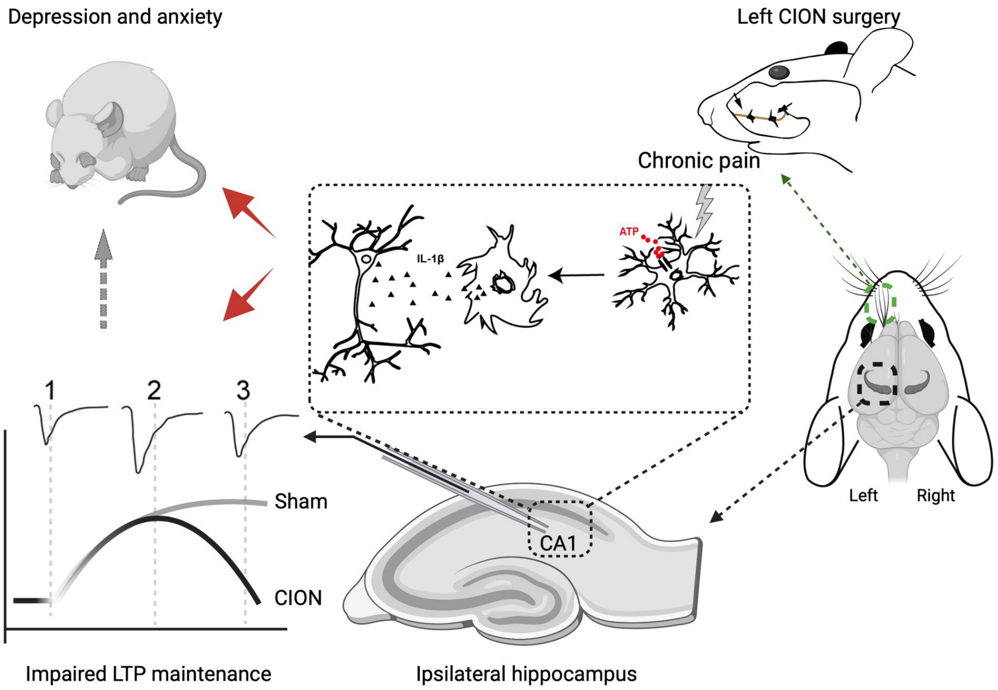


**Supplemental Figure 8. related to discussion, A schematic illustration of probable mechanisms for microglial activation in the ipsilateral hippocampal CA1 area impairing LTP and leading to anxiodepressive consequences of trigeminal neuralgia.** Increased extracellular ATP acts on P2X7 purinergic receptor on microglia and activates microglia. Activation of microglia results in increased IL-1 β release, leading to LTP impairment and anxiodepressive-like behaviors.
